# ACCLIMATION OF PHOTOSYNTHESIS TO THE ENVIRONMENT 1 regulates Photosystem II Supercomplex dynamics in response to light in *Chlamydomonas reinhardtii*

**DOI:** 10.1101/2020.02.26.966580

**Authors:** Marie Chazaux, Stefano Caffarri, Juliane Da Graça, Stephan Cuiné, Magali Floriani, Pawel Brzezowski, Gilles Peltier, Bernard Genty, Jean Alric, Xenie Johnson

## Abstract

Photosynthetic organisms require acclimation mechanisms to regulate photosynthesis in response to light conditions. Here, two mutant alleles of *ACCLIMATION OF PHOTOSYNTHESIS TO THE ENVIRONMENT 1* (*ape1*) have been characterized in *Chlamydomonas reinhardtii.* The *ape1* mutants are photosensitive and show PSII photoinhibition during high light acclimation or under high light stress. The *ape1* mutants retain more PSII super-complexes and have changes to thylakoid stacking relative to control strains during photosynthetic growth at different light intensities. The APE1 protein is found in all oxygenic phototrophs and encodes a 25 kDa thylakoid protein that interacts with the Photosystem II core complex as monomers, dimers and supercomplexes. We propose a model where APE1 bound to PSII supercomplexes releases core complexes and promotes PSII heterogeneity influencing the stacking of Chlamydomonas thylakoids. APE1 is a regulator in light acclimation and its function is to reduce over-excitation of PSII centres and avoid PSII photoinhibition to increase the resilience of photosynthesis to high light.

## Introduction

In plants, algae and cyanobacteria, photosynthetic electron transport is tightly linked to light capture and CO_2_ assimilation. Depending on light intensity and CO_2_ availability, either the light reactions or carbon metabolism will be limiting for photosynthesis (Farquhar et al., 1980). Balancing the two results in poising the electron carriers along the chain from the Photosystem II (PSII) electron acceptors (plastoquinones) to the Photosystem I (PSI) electron acceptors (Fd, FNR, NADP^+^). This energetic balance must be constantly tuned to environmental cues in order to avoid over-reduction of the photosynthetic apparatus and photo-damage. Referred to as photosynthetic acclimation, it involves both fast regulation of thylakoid proteins as well as slower, long-term changes to light harvesting and photosystem stoichiometry (for reviews see (Anderson et al., 1995; Erickson et al., 2015; Walters, 2005)).

PSII is the water splitting primary reaction component of the electron transport chain of oxygenic photosynthesis. It is a highly conserved multi-subunit complex composed of pigments and proteins which first evolved in cyanobacteria, the progenitor of chloroplasts (Caffarri et al., 2014). The PSII reaction centre (RC) is made up of D1 and D2 proteins that contain the co-factors required for charge separation, heme *b*_559_ (bound to PsbE and F), and the inner antenna of the core, CP43 and CP47 (Bricker and Ghanotakis, 1996). The PSII core complex includes the RC plus the oxygen evolving complex, PsbO, PsbP and PsbQ (Thornton et al., 2004). Additionally, around 20 other low MW proteins are observed in the homodimeric crystal structure and are required for PSII dimer or supercomplex stability (Caffarri et al., 2009; Umena et al., 2011). Many more proteins have been found to interact with PSII transiently in substochiometric quantities with precise functions to optimize oxygen evolution, electron transfer, PSII stability, protection or repair. These extrinsic proteins contribute to the flexibility and stability of PSII during periods of environmental variation or during biogenesis (Komenda and Sobotka, 2016; Plochinger et al., 2016; Shi et al., 2012). Efficient PSII regulation and repair is a necessity because PSII contains chlorophyll excited states that can react with locally produced oxygen. During repair, the turnover rate of D1 protein can be very high while electron transfer remains functional: this is achieved by dynamic changes involving the heterogeneity of PSII complexes (Guenther and Melis, 1990; Jarvi et al., 2015; Kirchhoff, 2019).

Large antenna complexes are required to efficiently capture the light energy for transfer to the PSII core. In all phototrophic eukaryotes, the light harvesting complex II (LHCII) proteins that are rich in chlorophyll *b* (chl *b*) and account for most of the pigments associated with PSII perform this role. The LHCII antennae attached to a PSII core form a PSII supercomplex (PSII SC) and this association can take different oligomeric forms and is determined by environmental conditions (Bielczynski et al., 2016; Shen et al., 2019; Su et al., 2017). Regulators LHCSR3 and PsbS, bind to the antenna of PSII SC and down regulate light harvesting by promoting non-photochemical quenching (NPQ), a safe dissipation of light energy as heat (Correa-Galvis et al., 2016a; Correa-Galvis et al., 2016b; Nawrocki et al., 2020; Semchonok et al., 2017; Tibiletti et al., 2016). State transitions change the functional antenna size of PSII whereby the phosphorylation status of the LHCII adapts antenna cross section of both PSI and PSII to the redox state of the electron transport chain (Dumas et al., 2016). Such quenching and photoprotection mechanisms are considered to be short-term acclimation processes (Erickson et al., 2015).

At the supramolecular level, PSII is embedded in thylakoid membranes of eukaryote phototrophs. Similar to plants, in unicellular green algae, PSII are mostly found in more stacked regions of the thylakoids while PSI are found in non-appressed lamellae and margins (Goodenough et al., 1969; Goodenough and Levine, 1969; Goodenough and Staehelin, 1971; Kouril et al., 2018). However, PSII is heterogeneous both in its distribution in the thylakoids and in its supramolecular organization. PSII SC are found in the grana stacks while PSII core complexes can also localize to margins and stromal lamellae (Albanese et al., 2016; Boekema et al., 1999; Danielsson et al., 2006; Drop et al., 2014; Koochak et al., 2019; Schwarz et al., 2018; Suorsa et al., 2015). These organizations are dynamic and thylakoids adapt their structure in response to light. In low light, thylakoids form stacks of highly dense membranes containing PSII SC, a structure favoring light capture (Polukhina et al., 2016; Rochaix, 2014). PSII forms semi-crystalline ordered arrays (Boekema et al., 2000) and high molecular mass assemblies containing numerous LHCII and forming megacomplexes across the stromal gap (Albanese et al., 2017; Albanese et al., 2016; Boekema et al., 2000; Wei et al., 2016). In high light, thylakoid membranes become less stacked and the number of layers decreases. This change can be as fast as a few minutes after the transition from low to high light (Rozak et al., 2002).

Destacking of thylakoid membranes is accompanied by PSII antenna size adjustments and changes to the density of SC, which corresponds to the long-term acclimated structure in response to the higher light intensity (Bielczynski et al., 2016; Kouril et al., 2013; Polukhina et al., 2016). Destacking allows for damaged PSII core complexes to disassemble into PSII core dimers, PSII core monomers and Repair Complex 47 (RC47: PSII core monomer lacking CP43) and facilitates migration towards the non-appressed lamellae and grana margins. Changes to stacking and reduction in antenna size of PSII prevents oxidative stress (Herbstová et al., 2012; Khatoon et al., 2009) and is also required for PSII repair (Jarvi et al., 2015; Theis and Schroda, 2016)

In eukaryote oxygenic phototrophs, the acclimation response to an increase in light intensity is defined by three major factors: a decrease in the relative abundance of thylakoid membrane to stroma, decrease in relative chlorophyll content and an increase to chlorophyll *a/b* ratio (Melis, 1996). With higher light the requirement for light harvesting is reduced, while the photosynthetic yield increases in line with electron transport and CO_2_ assimilation rates relative to the lower light intensity (Anderson et al., 1995). *Acclimation of Photosynthesis to the Environment 1* (*ape1*) allele was initially identified in a screen in *Arabidopsis thaliana* to identify mutants affected in the long-term light acclimation responses that are linked to increasing photosynthetic yield (Walters et al., 2003). At*ape1* did not increase PSII quantum yield after a shift to high light. During acclimation to strong light, the Chl *a/b* ratio increased in the wildtype, but not in At*ape1,* indicating that the high light response of increasing PSII core (rich in chl *a*) and decreasing LHCII antenna (rich in chl *b*) was active in the wildtype but affected in At*ape1* (Walters et al., 2003). This preliminary characterization pointed to a role for APE1 in an unknown but major light acclimation mechanism, but neither the mechanism nor the protein was further studied.

Using *Chlamydomonas reinhardtii* as a model organism, we present a characterization of high light acclimation and we demonstrate that the *ape1* mutant phenotype is deficient in this process. We have determined APE1 localization, interactions, and function in regulating light capture and photoprotection at the level of PSII. We propose that APE1 plays a role in the remodeling of PSII SC, thereby regulating electron transport for CO_2_ fixation and preventing PSII photoinhibition.

## Results

### *APE1* maintains photosynthetic growth in high light

In the context of deepening our knowledge on light acclimation processes, we isolated a mutant in Chlamydomonas with poor photosynthetic growth at high light affected for variable chlorophyll fluorescence under these conditions (Figure 1)(Figure 1A, B, C). Using a PCR-based technique we identified the left-flanking region of the insertion site of the antibiotic resistance cassette in the 3’UTR of the *Cre16.g665250* locus, annotated as *<u>A</u>cclimation of <u>P</u>hotosynthesis to the <u>E</u>nvironment <u>1</u>* (*APE1*, www.phytozome.jgi.doe.gov, v12.1.5). Sequencing of the insertion site showed that 1.5 copies of the *APHVIII* cassette are present (Supp. Figure 1). RT-PCR of the *Cre16.g665250* locus in this mutant showed presence of *APE1* transcript, not unexpected for an insertion in the 3’UTR (Figure 1D). This allele was designated *ape1-1* after successive outcrossings and backcrossings to a wild type 137cc strain.

**Figure 1.**
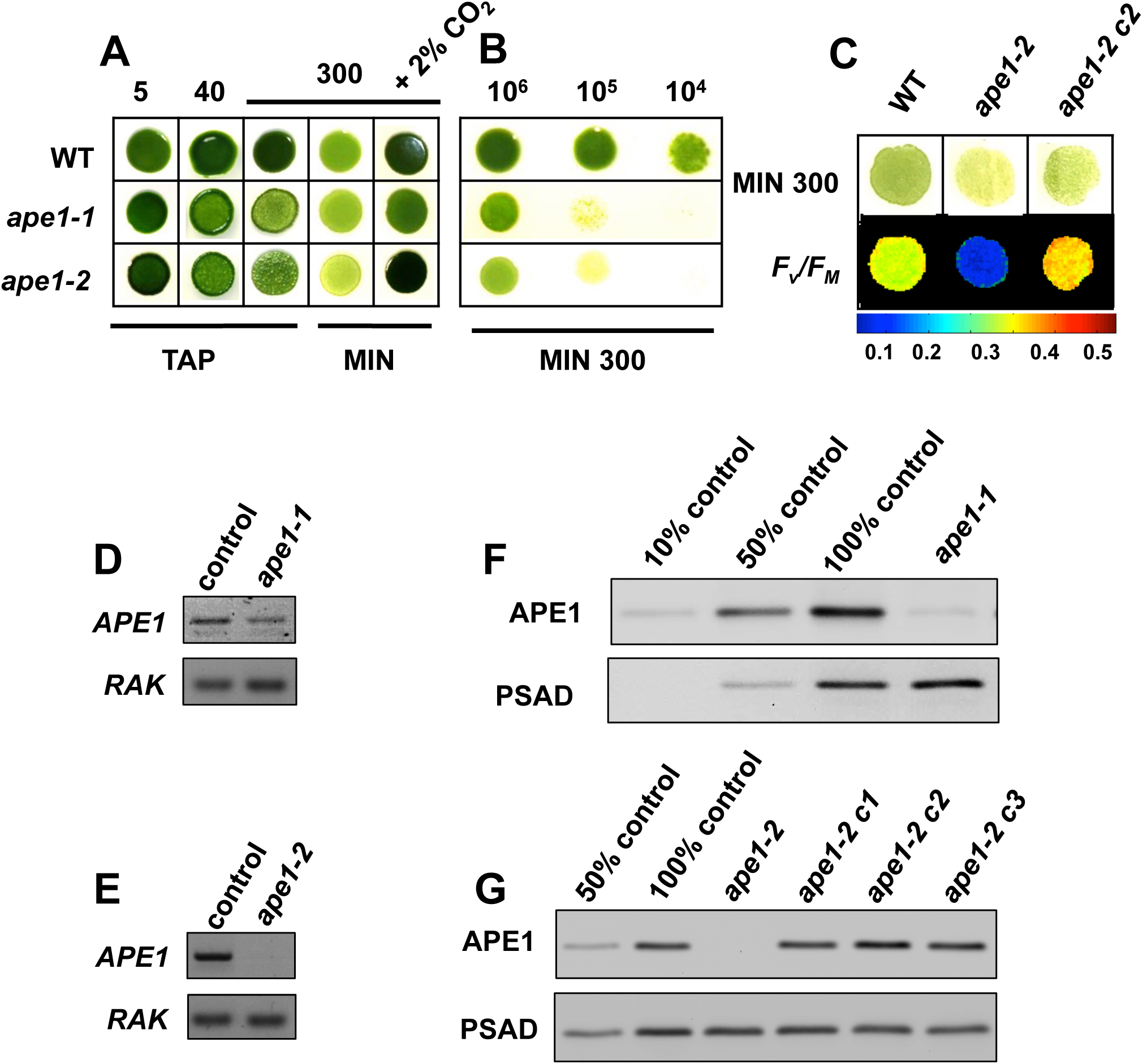
**Identification of *ape1* alleles that are interrupted in *Cre16.g665250* encoding APE1, a protein that contributes to optimal growth in high light.** A. Spot tests (10^6^ cells.mL^-1^) on rich media (TAP) versus minimal media (MIN) at different light intensities and with or without supplement of 2% CO_2_. B. Dilution series of cells grown at high light (300 μmol_photons_.m^-2^.s^-1^) on minimal media. C. Growth and maximum PSII quantum yield on minimal media grown at high light. The false color images give a range for the *F*_V_/*F*_M_ value. D. RT-PCR on control lines, *ape1-1* and E. *ape1-2* using *RAK* gene as a positive control and primers E and F (see Supplemental Figure 1) for *APE1*. F. APE1 immunoblot of total cell proteins of control line and wild type versus *ape1-1* and G. *ape1-2* and three complemented lines. PSAD was used as a loading control.

We also obtained a second allele from the CLiP library (Zhang et al., 2014), annotated as having an insertion in intron 4 of *Cre16.g665250*, and designated LMJ.RY0402.039504. The insertion site (Supp. Figure 1) was confirmed by PCR. RT-PCR analysis revealed no transcript for *APE1* (Figure 1E). The obtained mutation was successively crossed into our wildtype line and we named this allele *ape1-2*.

The two alleles *ape1-1* and *ape1-2* cells grew equally well as the wild type at 5 and 40 μmol_photons_.m^-2^ s^-1^ heterotrophically (see TAP in Figure 1A); on the contrary both *ape1-1* and *ape1-2* showed impaired growth in high light (at 300 μmol_photons_.m^-2^ s^-1^), especially on minimal medium (see the dilution series, Figure 1B). Addition of 2% CO_2_ in the growth chamber allowed the mutants to grow like the wild type. These observations seemed to assign to *APE1* a role in acclimation of oxygenic photosynthesis in response to light and CO_2_ availability.

Complementation of *ape1-2* line using the WT copy of APE1 under the control of its native promoter resulted in a complemented line (*ape1-2* c2) with improved phototrophic growth in high light compared to the *ape1-2* mutant (Figure 1C). Chlorophyll fluorescence imaging showed that the maximum quantum yield of PSII (*F*_V_/*F*_M_) was very low in the mutant (∼ 0.19) but it reached the WT (∼ 0.35) value in the complemented line (∼ 0.39).

Immunoblot analysis using the antibody against APE1 showed that this protein was reduced to around 5-10% of the wild type level in *ape1-1,* but it was absent from *ape1-2* (Figure 1F and G). The APE1 protein accumulation is restored to WT levels in three independent *ape1-2* complemented lines, which correlates with the improved growth characteristics (Figure 1C). The complemented line *ape1-2* c2 is referred to as *ape1-2*:APE1 in the experiments below.

### APE1 is a primordial thylakoid protein

The *APE1* gene model can be identified exclusively in oxygenic phototrophic organisms (Figure 2)(Figure 2A). APE1 has homologues in all cyanobacteria strains sequenced, including in *Gloeobacter violacelus* that lacks thylakoid membranes (Rexroth et al., 2011). It is however not found in any anoxygenic phototrophic species, bacteria containing only one of either type of reaction center (*Chloroflexus aurantiacus*, RCII or *Heliobacillus mobilis*, RCI). Supplemental Figure 2 shows representative species used for the hypothetical protein alignment used for the Cladogram. The predicted gene product shares 34-38% sequence identity with 17 homologs in these oxygenic phototrophs.

**Figure 2.**
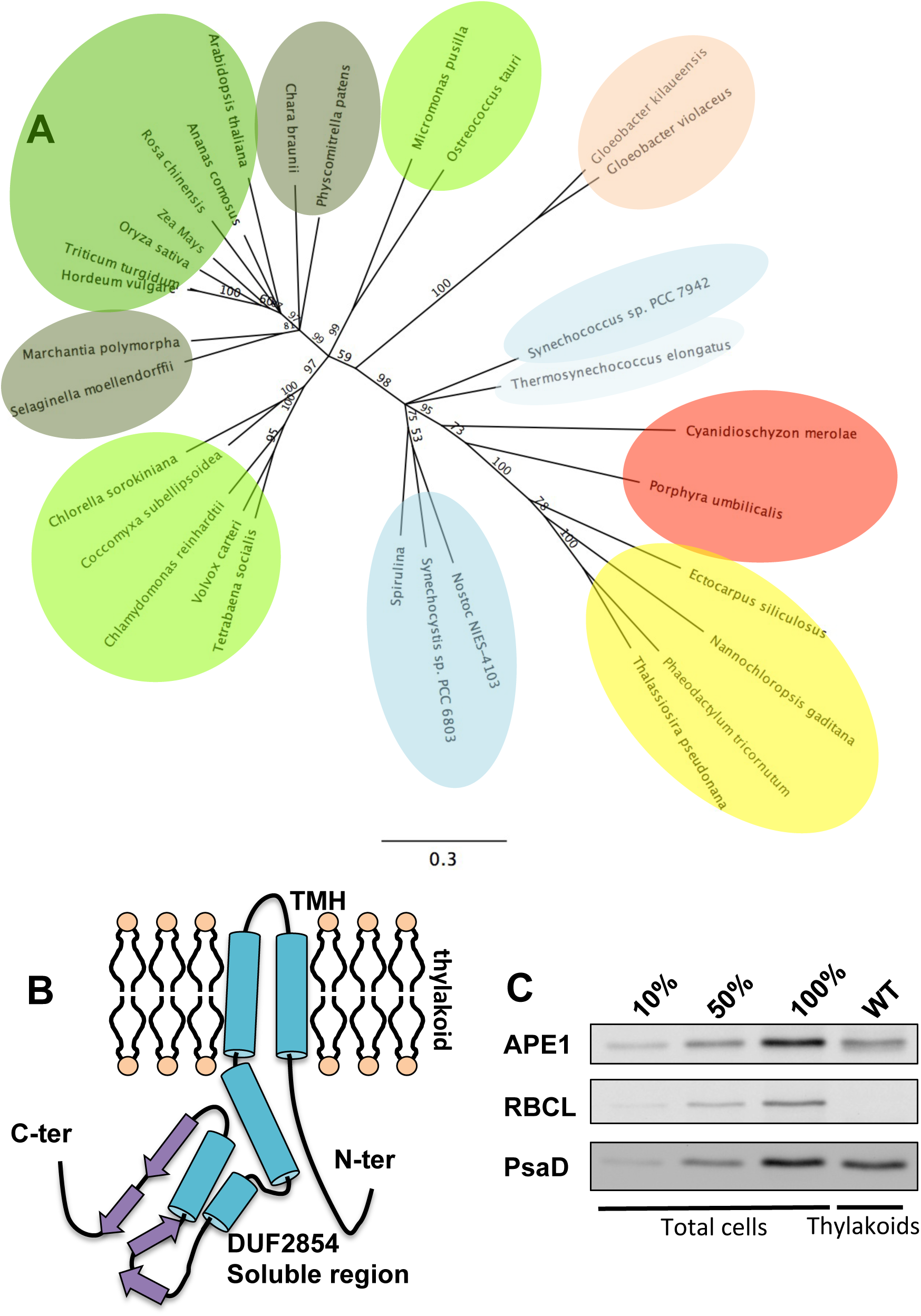
**APE1 is a thylakoid protein present in all oxygenic phototrophs. A.** Cladogram of APE1: Representative APE1 protein sequences from all clades of oxygenic photosynthetic species were identified by BLAST, alignment was done using Geneious (MUSCLE Alignment). The cladogram was constructed using a Jukes-Cantor genetic distance model neighbor-joining tree build method, unrooted, with Bootstrap consensus shown on branches. Colour codes represent phylum and distribution: green: land plants; army green: primitive plants; pale green: green algae; pale pink: primitive cyanobacteria; pale blue: hot spring cyanobacteria; blue: marine cyanobacteria; darker blue; fresh water cynaobacteria; pink: red algae; yellow: heterokont algae. **B.** Secondary structure model of the APE1 protein in the thylakoid lipid bilayer showing predicted (RaptorX) transmembrane and alpha helices (in blue) and beta-coil (in purple). **C.** Immunoblot of APE1 in total cells and thylakoids of the wild type. RBCL was used as a stromal control; PSAD was used as a thylakoid control.

The *APE1* gene of *C. reinhardtii* is predicted to encode a 25 kDa mature protein, modeled using RaptorX: http://raptorx.uchicago.edu/ (Kallberg et al., 2012) (Figure 2B) with two hydrophobic helices on the N-terminus (predicted as transmembrane helices by TMPred) and a soluble domain of unknown function (<u>D</u>omain of <u>Un</u>known <u>F</u>unction 2854) which constitutes the majority of the protein and is unique to APE1 (Supplemental Figure 2). The *C. reinhardtii* APE1 has a predicted chloroplast transit peptide of 41 amino acids (predicted by PredAlgo, giavap.genomes.ibpc.fr).

The APE1 antibody was used to determine APE1 protein localization. APE1 is present in the chloroplast, in the isolated thylakoid fraction, shown by immunoblot analysis. APE1 was present with thylakoid PSAD, while it was absent in the stromal fraction, where Rubisco accumulated (Figure 2C). APE1 is thus a chloroplast thylakoid protein with origins at the beginning of oxygenic photosynthesis.

### APE1 contributes to the accumulation of PSII proteins and LHCII antenna but is not required for biogenesis

To quantify possible differences in the accumulation of PS proteins, we analysed thylakoid proteins isolated from WT and *ape1-2* grown in low light heterotrophically. Proteins were separated using gradient denaturing gels and resolved using the highly sensitive SYPRO staining technique (Figure 3). Immunoblot analysis shows that APE1 cannot be separated from LHCII in this analysis (Figure 3A). PSII proteins D1 and D2 were quantified using Image J and found to accumulate less than the WT while LHCII proteins accumulate more than the WT (by approximately 20% in both cases). Other core and minor antenna appeared unaffected using this technique (Figure 3A). Immunoblot analysis shown for *ape1-1* grown phototrophically at low light (80 μmol_photons_.m^-2^.s^-1^) showed a similarly reduced PSII content for D1 and here the other PSII proteins tested were also reduced when compared against the wild type (Figure 3B). The accumulation of major photosynthetic complexes did not show major differences when respective protein content was compared. Photosynthetic parameters for cells grown under these conditions showed that the quantum yield of PSII was not significantly different in *ape1* from the control strains (Supp. Table 1).

**Figure 3.**
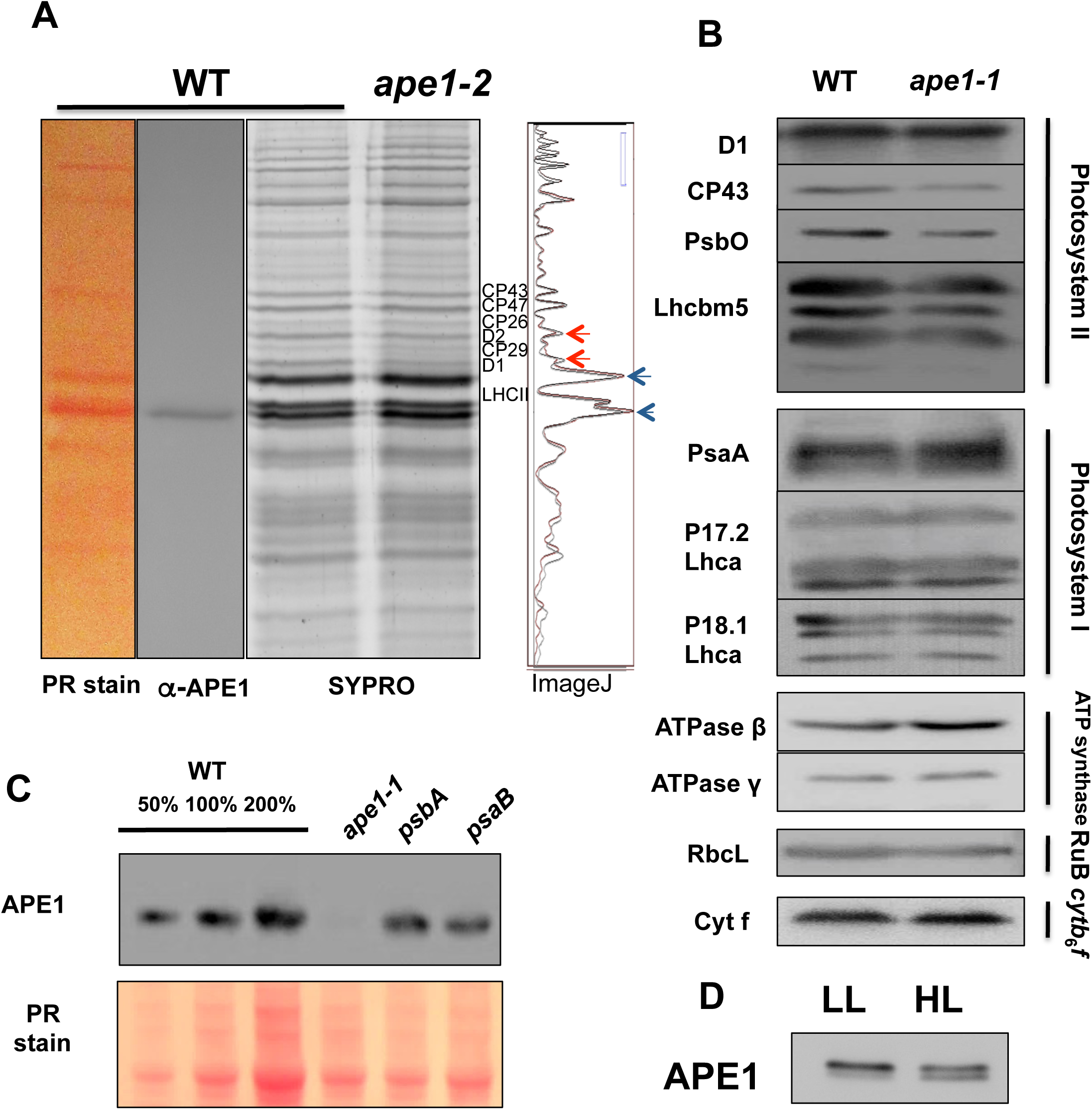
**APE1 contributes to PSII accumulation but is not required for photosystem biogenesis. A.** Thylakoid proteins were isolated from WT and *ape1-2* grown under low light (10 µmol_photons_.m^-2^.s^-1^) in heterotrophic conditions and separated using denaturing gels (15.5% 6M Urea) that were used for immunoblot of APE1 and for SYPRO staining. PSII proteins, minor antennae and LHCII were identified and quantified using image analysis (ImageJ). **B.** Immunoblot analysis of major photosynthetic complexes shown for wildtype and *ape1-1* grown at low light (80 µmol_photons_.m^-2^.s^-1^ under phototrophic conditions). **C.** Immunoblot analysis of APE1 accumulation in *ape1-1*, and mutants lacking PSII (*psbA*) and PSI (*psaB*). Ponceau Red was used as a loading control**. D.** Cells were sampled from photobioreactors at low (LL) and high light (HL) at the same cellular density and loaded onto denaturing gels. Western blot analysis was used to measure APE1 accumulation and bands were quantified using ImageJ.

When multi-subunits chloroplast complexes are not properly assembled, they are degraded, and when a dominant subunit of a complex is missing, the synthesis rate of the other subunits is reduced by the mechanism of “control by epistasy of synthesis” (Choquet et al., 2001). In *ape1-1*, APE1 barely accumulates (Figure 3C). In mutants devoid of PSII (*psbA*) and PSI (*psaB*), grown hetrotrophically, APE1 content was similar to the wild type. These results show that APE1 is not an intrinsic subunit of the photosystems and is not strictly required for their assembly.

The accumulation of the APE1 protein was also tested in both low and high light phototrophic culture conditions (Figure 3D). APE1 was quantified and found to accumulate to the same amount in both conditions. However as opposed to accumulation of APE1 in Figures 1, 2 and 3C showing protein samples from heterotrophically grown cells in low light, in figure 3D, APE1 migrates not as one band but as two bands and they accumulate in different quantities between low and high light phototrophic conditions. Taken together these results show that APE1 protein accumulates constitutively and may also be modified post-translationally depending on the media or the light intensity.

### APE1 interacts with PSII core complexes

The sequence analysis of APE1 and the low PSII quantum yield of *ape1* in high light suggested a functional link to PSII activity and we next performed experiments to identify protein-protein interactions with native photosynthetic complexes. Mild solubilization of purified thylakoid membranes with 0.5% digitonin/ 0.5% α-DM followed by separation of protein complexes by BN PAGE (4-16%) was followed by immunoblot analysis of D1 and PsaD to identify PSII and PSI complexes. It gave five bands for PSII: two SC forms, of which one also contains PSI, the PSII core dimers, the PSII core monomers and RC47 (PSII Reaction Center with CP47 but without CP43). Four bands of varying intensity were identified for PSI (Figure 4).

**Figure 4.**
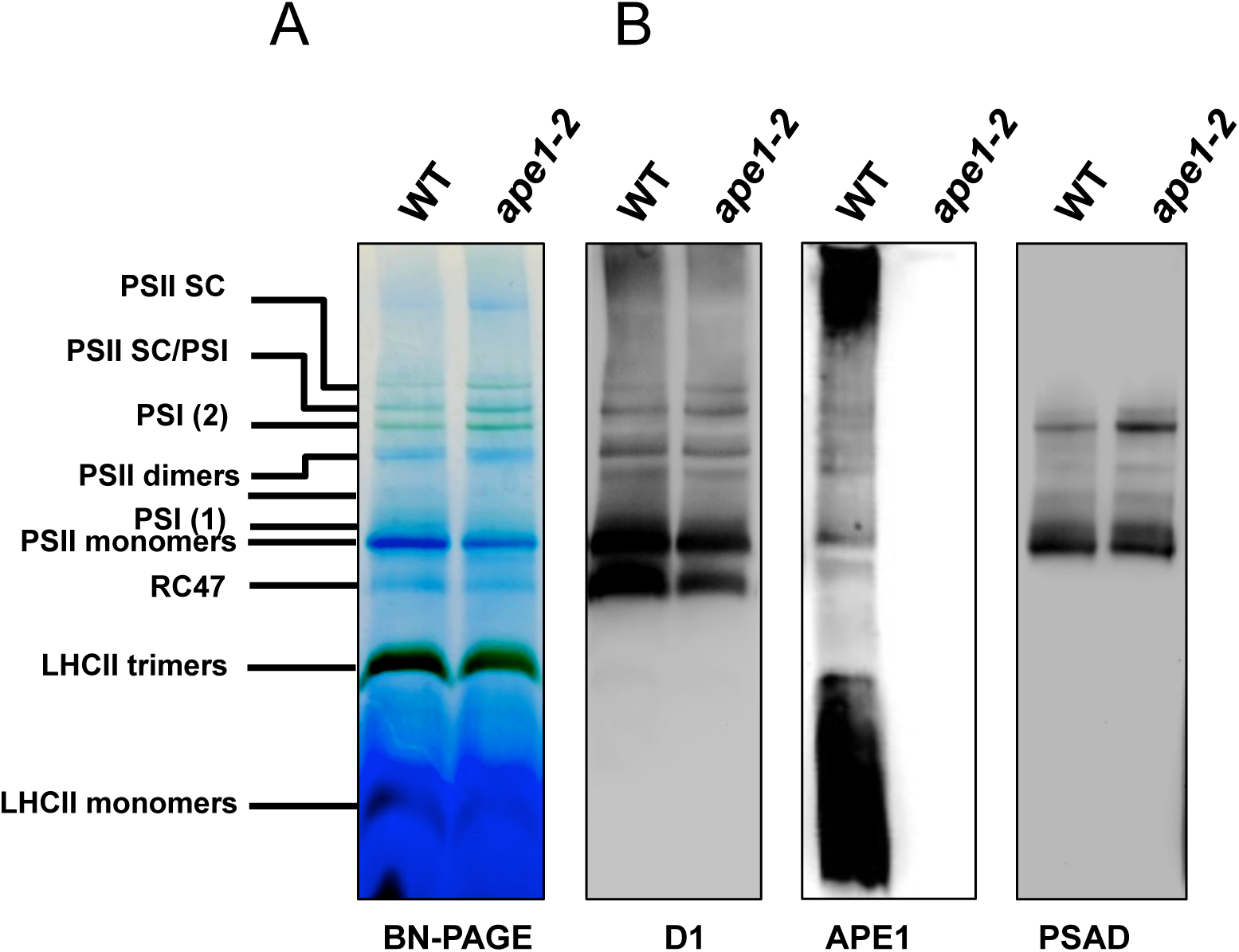
**APE1 affects the distribution of high molecular mass photosystem complexes and the APE1 protein co-migrates with PSII. A.** Analysis of native protein complex formation of WT and *ape1-2* cells grown phototrophically in low light. Thylakoid membrane proteins were solubilized using 0.5% digitonin/0.5% α-DM and separated by BN-PAGE (4-16%). Proteins were loaded on a per chlorophyll basis. **B.** Immunoblot analysis of the BN-PAGE in (A.) using antisera against D1 protein, APE1 and PSAD.

Immunoblot with APE1 antibody did not show any cross-immunoreaction against thylakoid proteins other than APE1 (see *ape1-2* lane, Figure 4B). APE1 migrated mostly with the higher molecular mass fractions (megacomplexes, aggregates or non-solubilized membrane fragments) as well as the low molecular weight fraction. In between these two extremes, defined bands were obtained at the same level of D1, showing that APE1 co-migrated with PSII SC, PSII dimers and monomers, but not the PSII repair fraction, RC47. Neither, did APE1 co-migrate with the trimeric antenna fraction. In BN PAGE (4-16%) there were at least two bands where PSI and PSII co-migrate or migrate very closely to each other.

Therefore, we tested for APE1 accumulation in complexes from mutants devoid of PSI (Δ*PsaB*) or PSII (Δ*PsbA*) and using BN PAGE (3-12%) and silver-stained 2D gels (Figure 5)(Supplemental figure 3). As in Figure 4B, APE1 was found below and above LHCII trimers, probably as aggregates. In Δ*PsaB*, APE1 was detected in two specific bands, PSII core monomers and dimers that are absent from the Δ*PsbA* lane. This result unambiguously shows that APE1 co-migrates with the PSII core, with monomers and dimers.

**Figure 5.**
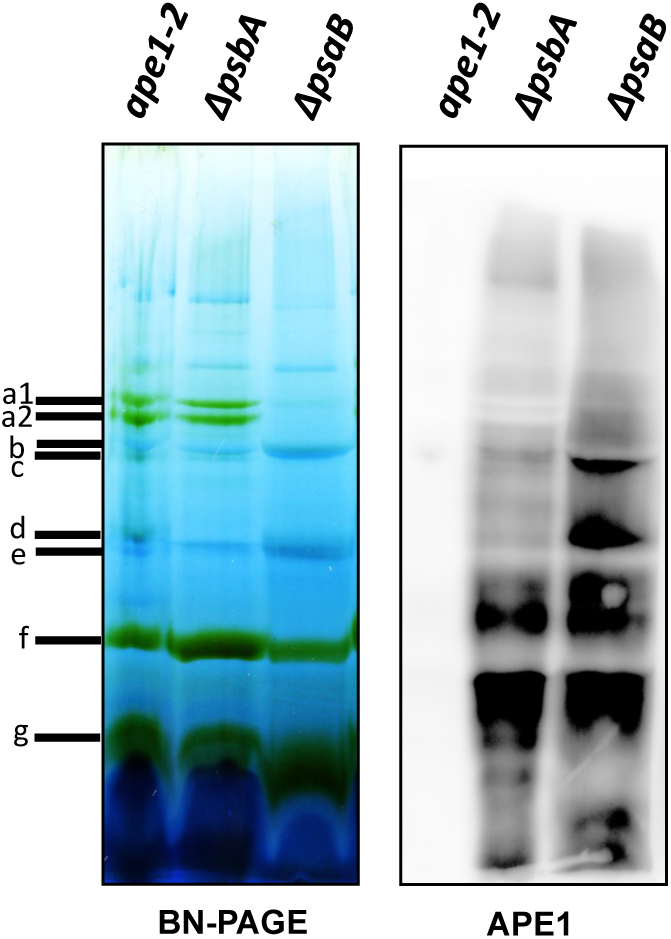
BN-PAGE (3-12%) analysis of *psbA* and *psaB* mutants grown heterotrophically in low light (10 µmol_photons_.m^-2^.s^-1^) with thylakoids isolated and treated under the same conditions as in Figure 4 and immunblot analysis using the APE1 antibody. Labelled bands were identified by excising the lanes and separation by SDS-PAGE in the second dimension : a1, PSI; a2, PSI; b, PSII dimers; c, PSI core; d, PSII monomers; e, cyt*b*_6_*f*; f, LHCII trimers; g, LHCII monomers.

To prove an interaction with PSII, protein crosslinking was tested using glutaraldehyde (GA) (Figure 6). Thylakoid membranes were solubilized in 1% α-DM and 1% Digitonin and loaded on sucrose gradients, with or without GA and subjected to ultracentrifugation. The sedimentation profile was not significantly changed whether the crosslinking agent was added or not (see Figure 6A), showing that, under these conditions, GA did not affect the fractionation into SC. Seven bands were collected and their absorbance was measured to control the content in photosynthetic complexes. All of the B5-B8 fractions contained PSII, the B5 fraction peaked at 675 nm with low emission for chl *b* region 630-660 nm, so we identify this as a fraction enriched in PSII core dimers. For B8 there is a relative increase in intensity around 630-660nm that we identified as LHCII of PSII SC (Figure 6B). The different fractions were then analyzed by denaturing SDS-PAGE. Except B1 (free pigments and small polypeptides) all fractions contained LHCII polypeptides, LhcbM1-8, (Drop et al., 2014) migrating between 25 and 30 kDa whether the cross-linking agent was present or not (this can be seen by comparing B7- and B7+ in Figure 6C). This shows that GA added to the gradient did not unspecifically cross-link all proteins into complexes, as also observed in the original protocol that was developed to strongly limit intercomplex crosslinking (Stark, 2010). B2+ and B3+ fractions did not contain high MW polypeptides whereas B5+ to B8+ contained large amounts of cross-linked polypeptides in the top part of the gel.

**Figure 6.**
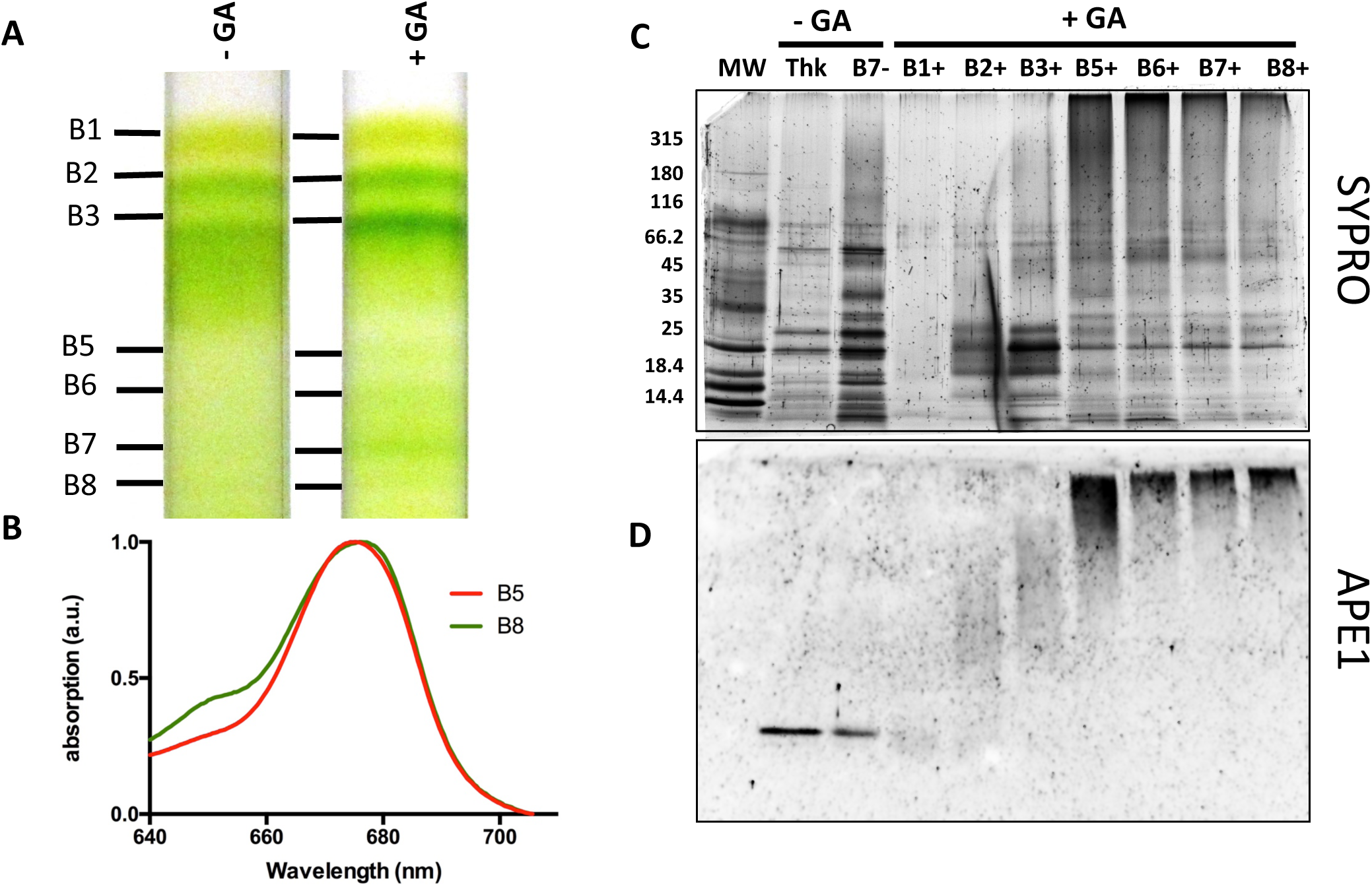
**Crosslinking of native thylakoid complexes reveals the interaction between APE1 and PSII core. A.** Thylakoid membranes extracted from wildtype cells grown phototrophically in low light were solubilized in 1% α- DM and 1% Digitonin, and loaded on sucrose gradients, with or without addition of glutaraldehyde (GA) as a crosslinker and subjected to ultracentrifugation. Labeled bands were isolated by syringe and these fractions were identified by their absorption spectra. **B.** absorption spectra at room temperature of the fractions B5 and fraction B8 from sucrose gradients. The spectra were normalised to maxima and minima in the 640-710 nm region. **C.** The different fractions with or without addition of glutaraldehyde (GA) shown in (A.) were analyzed by denaturing SDS-PAGE and stained with SYPRO, alongside the molecular weight marker (MW) and a sample of the non-treated thylakoid membranes (Thk). **D.** Immunoblot analysis of the gel shown in (B.) decorated with the APE1 antibody.

The immunoblot of the SDS-PAGE gel decorated with APE1 antibody in figure 6D shows that all cross-linked bands containing PSII (B5+ to B8+) contained the APE1 polypeptide in the upper part of the gel. B5+ lane showed a specific cross-linking of APE1 with the PSII core and only in the high MW part of the gel while it appeared as a single polypeptide in the non-treated thylakoid samples (Thk). APE1 does not migrate alone or in aggregates when the crosslinker is added showing that when protein-protein interactions are stabilized with GA, APE1 is indeed a PSII protein and not a free protein in the membrane.

### PSII is more prone to photoinhibition in *ape1*

Light sensitivity and a decreased maximum quantum efficiency of PSII in high light were observed in *ape1-2* (Figure 1 and Photosynthetic parameters in Supplemental Table 1), this appeared to be photoinhibition of PSII and defined as destruction of PSII centers occurring at a faster rate than they can be repaired (Adir et al., 2003). To determine whether lower PSII activity in the mutant was due to faster photodamage or due to impaired PSII repair and/or *de novo* synthesis, we applied a photoinhibitory light treatment of 1800 μmol_photons_.m^-2^.s^-1^ for one hour, with recovery in low light (Figure 7). PSII maximum quantum efficiency and D1 accumulation were monitored (Figure 7A and B). To differentiate between PSII degradation rate and PSII synthesis rate we also performed the experiment in the presence of protein translation inhibitors (lincomycin, LC and chloramphenicol, CAP).

**Figure 7.**
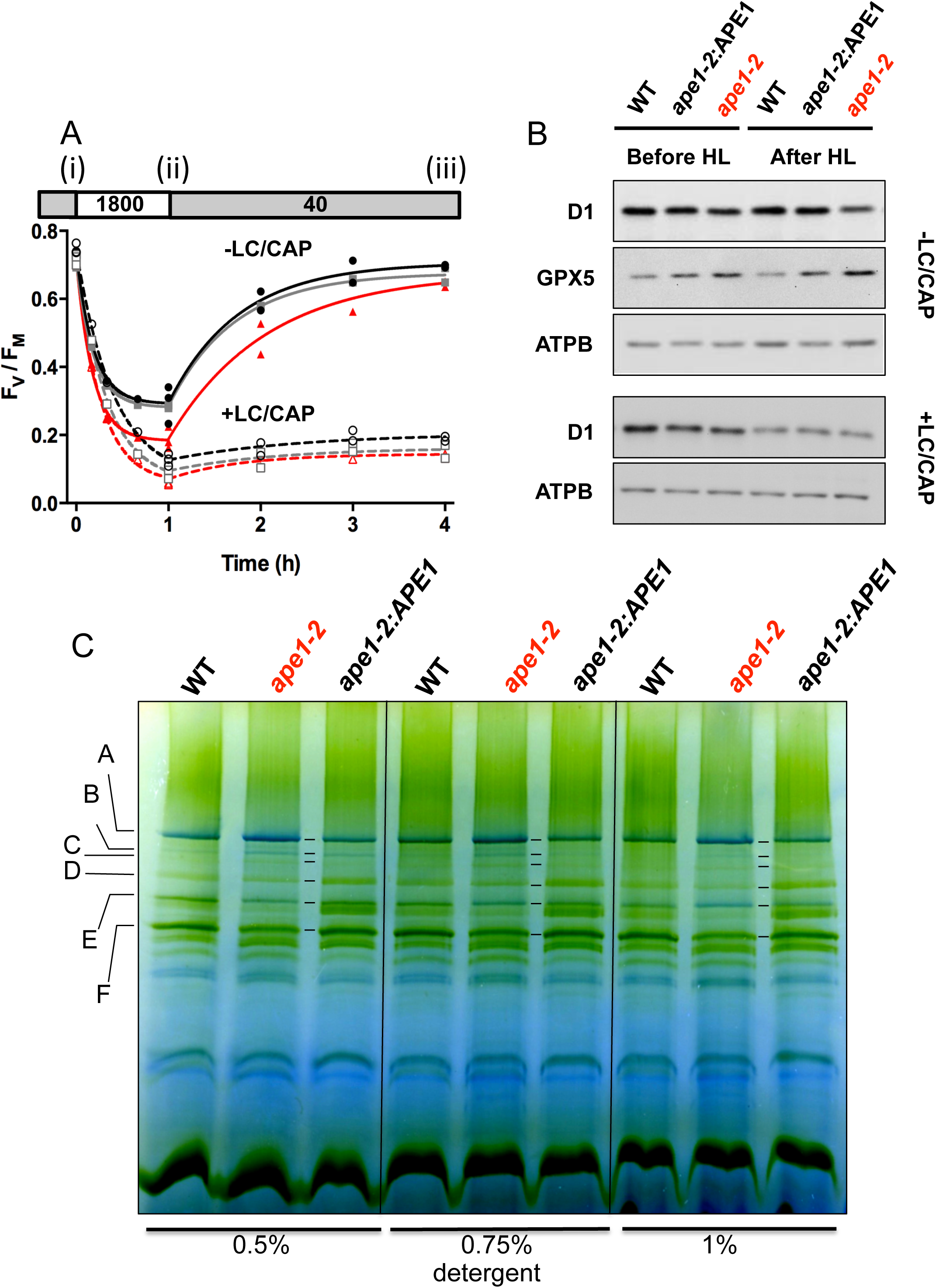
**APE1 protects against photoinhibition. A.** The wild type (black solid line), *ape1-2* complemented (grey solid line) and *ape1-2* (red solid line) strains in TAP media at the same cellular density were placed at (i) 40 µmol_photons_.m^-2^.s^-1^ for 60 min, (ii) photoinhibited (PI) for 60 min at 1,800 µmol_photons_.m^-2^.s^-1^, and then (iii) allowed to recover at 40 µmol_photons_.m^-2^.s^-1^. PSII fluorescence was measured by pulse amplitude-modulated fluorometry. Lincomycin and chloramphenicol were added (+LC/CAP) to inhibit translation of chloroplastic proteins (dashed lines). Two replicates are shown and curves were fitted using a single exponential. **B.** Immunoblot analysis of D1 and GPX5 (monitoring ROS production) accumulation during the experiment shown in (A.), before and after the high light treatment, both without and with inhibitors of protein translation. ATPB was used as a loading control. **C.** Cells from *ape1-2*, WT and *ape1-2*:APE1 were grown phototrophically in batch cultures at ambient CO_2_ at low light and then subjected to high light (300 μmol_photons_.m^-2^.s^-1^) overnight (16 hours). Thylakoids were isolated and the concentration of the solubilizing agents α-DM and digitonin were varied from 0.5 to 1 % for the same concentration of chlorophyll per sample and separated by BN PAGE (3-12%). The complexes showing the major differences between the *ape1-2* and the control lines were identified in *ape1-2* and represented by the letters: . A. PSII SC, PSI, mitoATPsynthase, B. PSII SC, mitoATPsynthase, C. PSII SC, mitoATPsynthase, D. PSII SC, mitoATPsynthase, E. PSII core dimers, LHCII, PSI, F. PSI

As expected, the presence of translation inhibitors (+LC/CAP) prevented the recovery of *F*_V_/*F*_M_ after photoinhibition (1 to 4 hours in Figure 7A). This was due to the block of PSII turnover and D1 synthesis, observed as a decrease in D1 accumulation in the bottom panel of Figure 7B. In the absence of translation inhibitors (-LC/CAP), a smaller decrease in D1 content was observed after an hour (Figure 7B), suggesting that in this instance D1 synthesis partially compensated D1 degradation. The same was also observed on the amplitude of *F*_V_/*F*_M_, consistently greater at 1 hour, showing that PSII was turning over in -LC/CAP conditions (Figure 7A).

The decrease in photosynthetic parameters (Fv/Fm) was paralleled by a decrease in D1, more pronounced in the mutant than in the reference strains (Figure 7B). Since the recovery rate in the mutant was similar to that of the wild type and complemented strain, decrease of D1 in *ape1-2* cells was more due to a faster PSII photoinhibition than a slower PSII repair.

The detoxification enzyme, Glutathionine Peroxidase 5 (GPX5) is a marker for singlet oxygen and PSII dysfunction (Fischer et al., 2009; Roach et al., 2017). GPX5 levels were higher in the mutant already in low light, and its content increased after the photoinhibitory treatment (Figure 7B). This result suggests that a lack of APE1 causes ROS production that elicits a systemic response that we witness by GPX5 accumulation in the mutant, and that the ROS produced is likely to be singlet oxygen issue of PSII.

We next examined the formation of high molecular mass complexes after a high light treatment. We tested different detergent concentrations and found that the dose did not significantly change the fractionation of complexes which remained stable for each strain (Figure 7C). Notably, the absence or presence of APE1 had a major impact on the pattern of high molecular mass complexes (A-F in Figure 7 and supplemental figure 4). The heaviest band, A, that appears blue and of greater abundance in *ape1-2* is composed of PSII SC, PSI and mitochondrial H^+^ATPsynthase (the latter is a contaminant from the thylakoid isolation common in cells grown under these conditions (Rexroth et al., 2003)). The bands B-E are more delineated but less green, suggesting bleaching, compared to the controls. The PSI band, F, is less abundant, suggesting a shift to a higher molecular mass band A in comparison to controls. Importantly, under these conditions, *ape1-2* retains PSII dimeric cores but is lacking the PSII monomeric core and RC47 complexes with an absence of CP43, CP47, D1 and D2 proteins at the expected position in the 2D gel (Supplemental Figure 4).

### PSII antenna size, heterogeneity (α- and β-centers) and connectivity between PSII centres

Differences in PSII photoinhibition linked to changes to PSII SC observed in BN-PAGE (Figure 7) led us to test *in vivo* the effective antennae size of PSII in *ape1* under physiological conditions. During a pulse of saturating light Q_A_ reduction rate is faster than Q_A_ to Q_B_ electron transfer and the variable chlorophyll *a* fluorescence increases from F_0_ to F_M_ and is commonly referred to as an OJIP induction. The first phase of this fluorescence rise (OJ phase ∼1 ms) reflects PSII antenna size, the faster the rise the larger the functional antenna size (Dinc et al., 2012) (Figure 8). We used a similar treatment as to that shown for the photoinhibition experiment (Figure 7A) and found that *ape1-2* PSII antenna size was slightly increased in comparison to the controls for conditions (i) and (iii) (Table I and Figure 8A). This suggests that *ape1-2* tends towards a larger PSII antenna than both the control lines becoming significantly different after the high light treatment (Table I).

**Figure 8.**
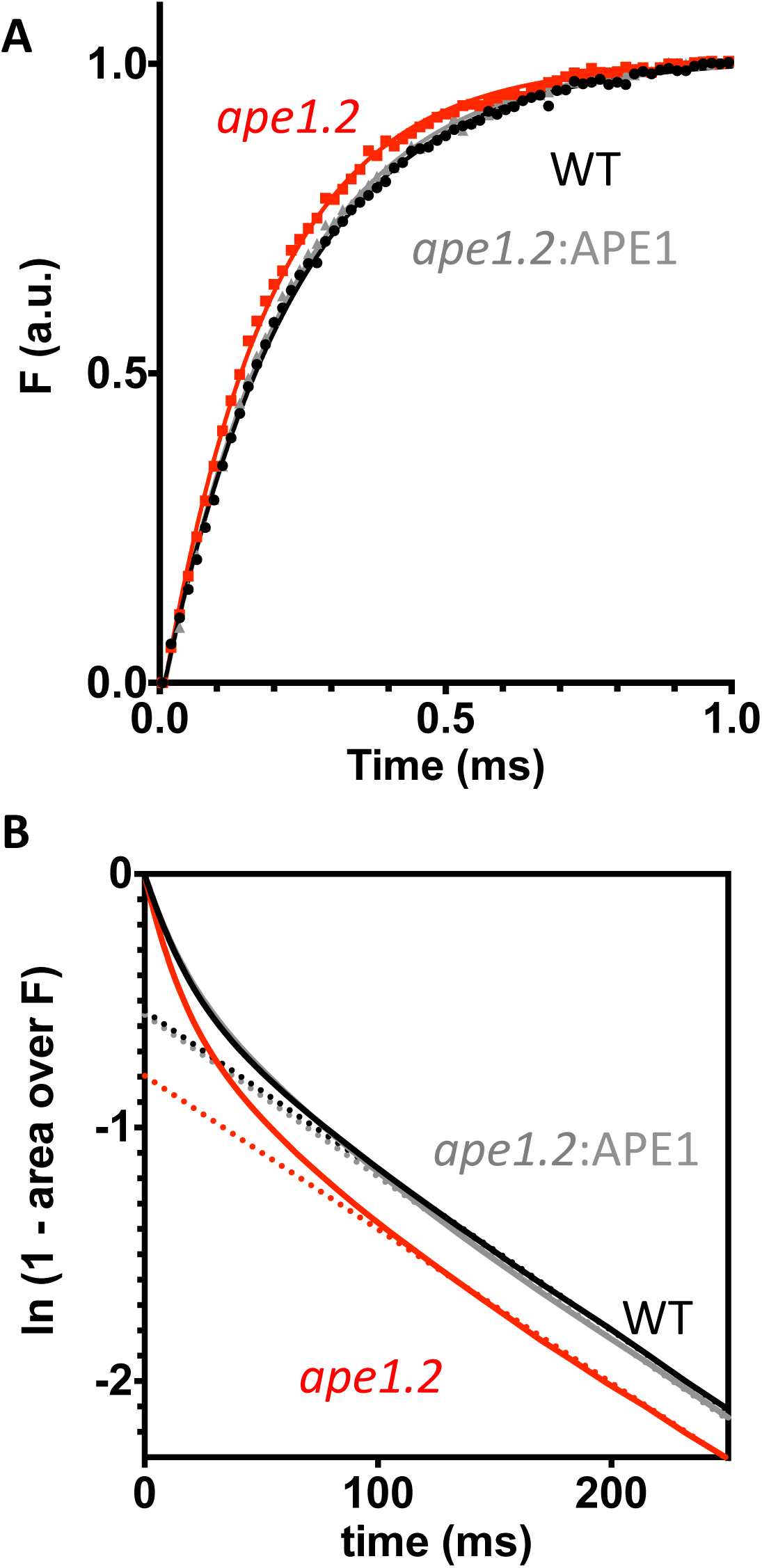
**Differences in effective antenna size can be measured *in vivo*** between wild type (black), *ape1-2* (red) and *ape1-2*: APE1 (grey) **A.** A typical fluorescence rise (F) for intact cells at room temperature pre-treated with high light (500 µmol photons.m^-2^.s^-1^) with recovery in low light (40 µmol photons.m^-2^.s^-1^) as in (iii) Figure 7. Actinic light was 8000 µmol photons.m^-2^.s^-1^. Data is normalized with F = 0-1 **B.** Estimation of the relative fraction of α-centers, i.e. large antenna size Photosystem II. The bi-phasic Q_A_ reduction rate as the time-dependent complementary area over the fluorescence rise, shown in logarithmic scale. Dotted lines intercept the ordinate axis at t = 0, yielding 55% α-centres for *ape1-2* and 42% for wild type and *ape1-2*:APE1. The experiements shown are representative of n>3 showing similar trends.

**Table I.** Fluorescence half rise times are shown for *ape1-2* and control lines before the light treatment, (i) and after (iii) using mean values (n=3) with SEM. * denotes that *ape1-2* was p < 0.05 significantly different from the controls.

|  | condition (i) |  |  | condition (iii) |  |  |
| --- | --- | --- | --- | --- | --- | --- |
|  | Low light |  |  | Low light acclimated after high light |  |  |
|  | WT | *ape1-2* | *ape1-2*:APE1 | WT | *ape1-2* | *ape1-2*:APE1 |
| F t<sub>1/2</sub> (ms) | 0.199±0.017 | 0.171±0.013 | 0.189±0.011 | 0.214±0.018 | 0.175±0.005* | 0.225±0.009 |

We questioned whether APE1 was affecting the excitonic connectivity of PSII centres, following the method of (Cuni et al., 2004). There were only small differences in the connectivity parameter J between *ape1-2* and WT prior to light treatment (i) (J_wt_ = 3.2±0.3 and J*_ape1-2_* = 4.1±0.5) and after recovery from the light treatment (iii) (J_wt_ = 3.1±0.2 and J*_ape1-2_* = 3.8±0.4). A significant deviation was observed at the light treatment (ii) in both strains (J_wt_ = 1.4±0.1 and J*_ape1-2_* = 1.5±0.1). This may be due to a smaller fraction of active PSIIs. During photoinhibition, the inactivation of PSII decreases the number of exciton traps as well as the probability of an exciton to hop from center to center. An absence of APE1 did not have an affect on this outcome.

Heterogeneity of PSII antennae was also tested in the presence of DCMU under non-saturating light. Under such conditions, the fluorescence rise is multiphasic. Analysis according to (Melis and Homann, 1976) showed that two phases are observed, one representing the α-centres or large antenna PSII and the other β, or low antenna PSII; no significant differences were observed for conditions (i) but in condition (iii), *ape1-2* had a larger amplitude of the fast phase α (∼55%) than the control lines (∼42%) suggesting a greater proportion of large antenna PSII SC (Figure 8B).

### APE1 changes fractionation profiles of PSII SC during light acclimation

To further characterize the different PSII SC partitioning, *ape1-2* and wild type were grown phototrophically in low and high light at ambient CO_2_ in turbidostats. Photosynthetic parameters were monitored (Supplemental table 1). At both light regimes, samples were collected after an acclimated state was reached (around 4 days). Using these conditions, six bands could be identified that contained PSII: PSII SC III, PSII SC II, a band containing PSII and PSI (PSII SC I/PSI), PSII core dimers, PSII core, and RC47 (Figure 9)(labeled in Figure 9A and 9C); and 5 containing PSI (Identified in Supplemental Figure 4).

**Figure 9.**
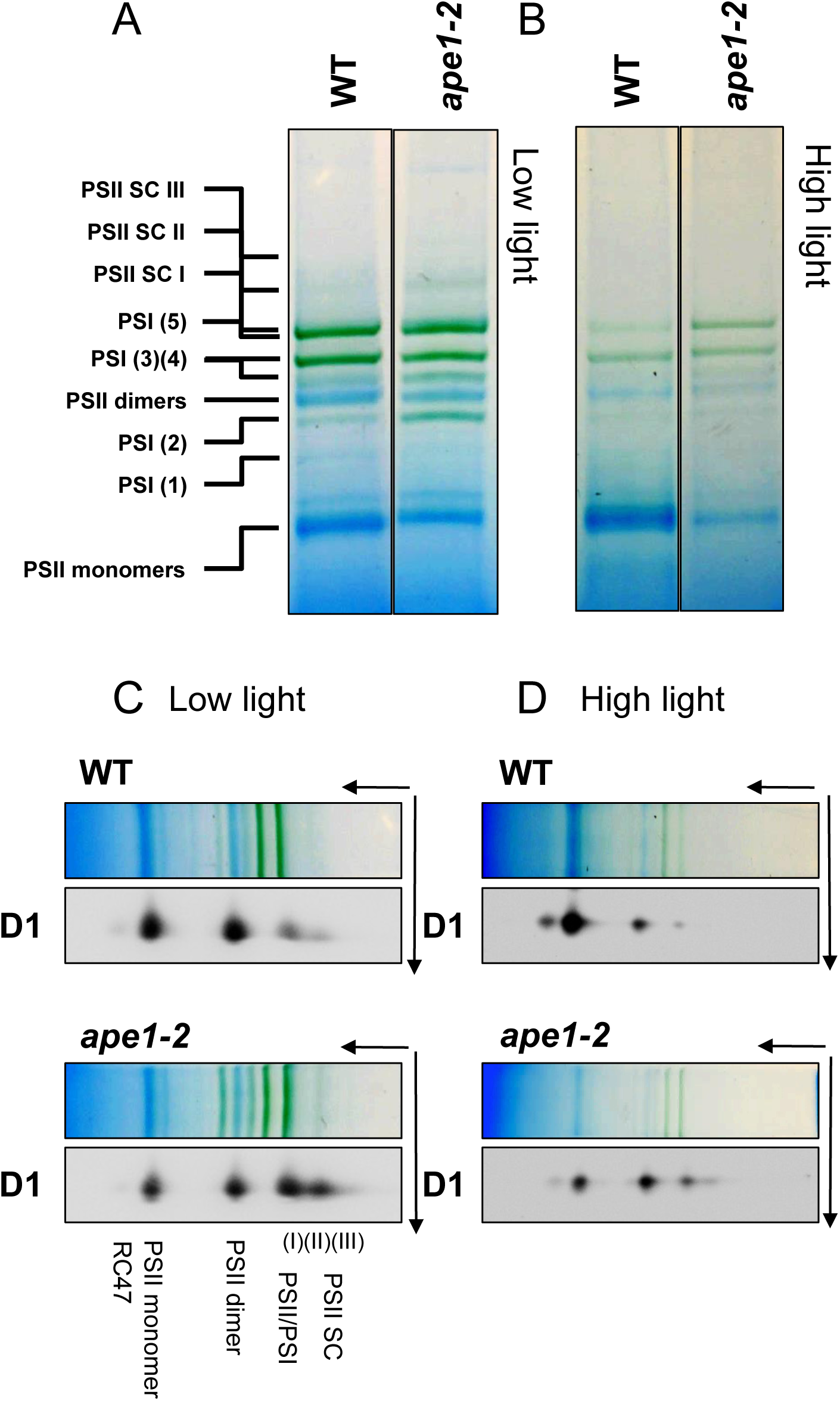
**APE1 participates in the mobility of PSII complexes towards core monomers in response to an acclimation to high light A.** BN-PAGE analysis of thylakoid proteins from wild type and *ape1-2* cells acclimated to low light and **B.** acclimated to high light. Thylakoid membrane proteins were solubilized using 1% β-DM and Digitonin and separated by BN-PAGE (4-16%). Proteins were loaded on a per chlorophyll basis. **C.** and **D.** The second dimension analysis was performed by excising and dentauring the BN-PAGE lane and proteins were separated by SDS-PAGE followed by immunoblotting using antisera against PSII D1 protein.

In low light conditions, in the wild type, PSII was mostly identified as dimeric and monomeric core complexes (Image J relative quantification of D1 from the immunoblot measured: 23% PSII SC; 37% dimeric and 37% monomeric cores; 3% RC47) (Figures 9A and 9C). The *ape1-2* profile was different; PSII SC were more highly represented, observed by D1 signal in the western blot (ImageJ relative quantification of D1 from the immunoblot measured 55% PSII SC; 22% dimeric and 22% monomeric cores; 1% RC47). The PSI profile was also different in *ape1-2*, with a higher abundance of higher molecular mass forms. These features regarding both PSII and PSI confirmed what was observed using milder solubilization.

The high light acclimation treatment (Figure 9B and 9D and silver stained gels in Supplemental Figure 4) resulted in changes to the PSII organization profile in the wild type. The ratio of high to low molecular mass assemblies decreased, where the signal for PSII SC, PSII SC I/PSI and PSII core dimers diminished in favor of PSII core monomers. PSI organization was also changed, with a reduction in the high MW bands. In *ape1-2*, the profile of PSII was less flexible: the same forms of PSII, from RC47 until PSII SC II, were still detectable. This led to a higher ratio of PSII SC to PSII core monomers in the absence of APE1 (Figure 9). Interestingly and despite significant photoinhibtion of *ape1-2*, CO_2_ fixation was not affected in comparison to the control strains (Supplemental table I).

### Thylakoid structure is altered in *ape1*

Thylakoid membrane structure is determined by a number of factors including photosystem composition, distribution, LHCII interactions and modifications (Kirchhoff, 2019). Interaction of APE1 with PS II, and changes in photosystem oligomeric composition could be linked to modifications at the level of the thylakoid membrane. To verify this, *ape1-2* and wild type cells were grown in phototrophic conditions in low light and ambient CO_2_ and the thylakoid membranes were resolved by transmission electron microscopy (Figure 10).

**Figure 10.**
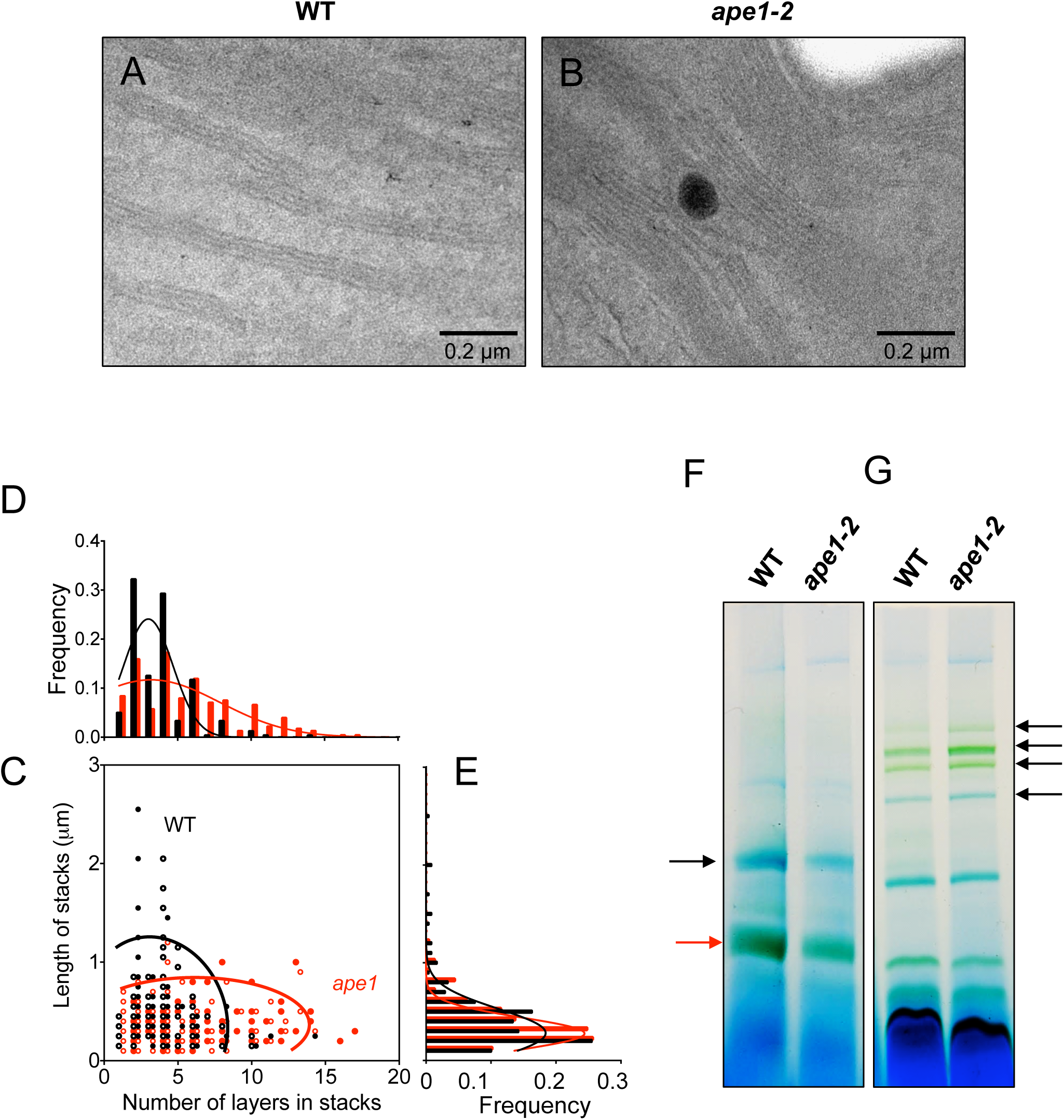
***ape1* has shorter thylakoid stacks composed of more layers, its thylakoids are enriched in grana cores in comparison to end membranes and stromal lamellae.** Transmission Electron Microscope images of thylakoids of **A.** wild type and B. *ape1-2* from cells acclimated phototrophically in turbidostats at low light and ambient CO_2_. C. Comparison of the stacking of thylakoid membranes between *ape1-2* and wild type. The length of a stack is represented against the number of layers as a scatter plot with 95% confidence ellipses calculated with Mathlab. **D.** and **E.** Data are projected into a single dimension giving the distribution of the number of layers and the length of the stacks with bell curves. Analysis is representative of 450 stacks and 85 TEM images. **F.** Differential solubilization of thylakoids of cells grown phototrophically at low light in batch cultures. Thylakoid membrane proteins were first solubilized using 1% digitonin to isolate end membranes and stromal lamellae and separated by BN-PAGE. Red arrow LHCII, balck arrow PSII **G.** The remaining pellet containing mostly stacked regions was then solubilized with 1% β-DM and separated by BN-PAGE. Arrows show differential accumulation of PSII enriched bands between the mutant and wild type in the two solubilized fractions.

The wild type appeared to have more organized thylakoid membranes as opposed to *ape1-2*, where thylakoids showed disorganized or uneven margins (Figures 10A and B and Supplemental Figure 5). Quantitative analysis (Figure 10C) showed that the mutant thylakoid stacks had more layers, often more than 7; against 3 - 4 on average in the wild type (Figure 10D). *ape1-2* stacks were shorter, always less than 1 μm, while the wild type showed membranes appressed over > 1 μm (Figure 10E). Statistical analysis showed that the differences observed were significant.

Calculation using the parameters found in Figure 10C and in (Engel et al., 2015) showed that the *ape1-2* mutant shows a 50% increase in the ratio of grana height to diameter compared to the wild type, and a 40% increase in the area of appressed to unappressed membranes (Supplemental Table 2). This is due to a larger stacked area and to a decreased end membrane surface, even though the area of the margins is about 20% higher in the mutant.

Differential solubilization of thylakoids of the wild type and *ape1-2* was performed to correlate changes in thylakoid structure with the localization of the photosystems (Figures 10F and 10G). We used digitonin to preferentially solubilize end membranes/stroma lamellae of thylakoids and then solubilized the remaining pellet (assumedly mostly stacked regions/grana) with β-DM and these were separated by BN PAGE (4-16%). We observed a lower yield on a per chlorophyll basis for digitonin solubilization in *ape1-2*, suggesting less LHCII (red arrow) and a reduction in low molecular mass PSII (black arrow), expected to accumulate preferentially in the end membranes and stromal lamellae regions as seen in the wild type (Figure 10F). The subsequent treatment of *ape1-2* with β-DM showed enrichment in high molecular mass oligomeric forms of PSII (black arrows), in the more stacked regions of the membranes (Figure 10G).

## Discussion

Structure and function of the photosystems and their arrangement in the thylakoid membrane are constantly adjusting to changes in their environment allowing them to maintain performance and to cope with stress. Modulating PS II oligomeric composition and antenna size in response to high light is a key parameter for light acclimation. The sum of acclimation mechanisms that an organism has at its disposal contributes to the fitness of a species and understanding these is fundamental to our comprehension of ecology and evolution, and to the future improvement of CO_2_ capture and utilization by photosynthetic organisms.

Proteomic studies in Chlamydmonas (Terashima et al., 2011), maize (Majeran et al., 2008) and Arabidopsis (Myouga et al., 2018; Tomizioli et al., 2014) have listed APE1 amongst the peptides found in thylakoid membranes. Western blot analysis on fractionated chloroplasts demonstrated that APE1 is an integral membrane protein, bound to the thylakoids (Figure 2). APE1 belongs to the GreenCut, the pool of gene models conserved across plants and green algae and this protein is absent from non-photosynthetic organisms (Merchant et al., 2007). More specifically, APE1 is part of the ‘PlastidCut + cyanobacteria’ (Heinnickel and Grossman, 2013). Comparative genomics of cyanobacteria showed that APE1 is part of their core genome, encompassing only 63 gene models shared with all oxygenic photosynthetic organisms (Mulkidjanian et al., 2006). The other conserved regulatory proteins in this group include GUN4 and THF1. A more recent analysis (Beck et al., 2018) compared the genome of 77 cyanobacterial species including the oceanic nitrogen-fixing cyanobacterium UCYN-A and *cyanobacterium endosymbiont of Epithemia turgida EtS* that have lost PSII and the genes for carbon fixation (Zehr et al., 2008). From the cross analysis of these different data sets we identified co-occurrence of APE1 with 7 PSII core subunits, that is, at the origins of oxygenic photosynthesis (Supplemental Table 3). This very early conservation may suggest that we have identified APE1 in the regulatory role that permits PSII heterogeneity and increases fitness and flexibility of the photosynthetic apparatus across all oxygenic photosynthetic species. However, the *ape1* phenotype is only observed under high light that suggests it became less important during evolution, as the repertoire of PSII repair and regulatory pathways has expanded over time. Rather than being a sole regulator it is probably now integral to a set of pathways involved in light acclimation (such proteins as CURT and STT7/8 could be considered to fall within the same category) that together optimize thylakoid structures in line with light conditions (Pribil et al., 2014; Pribil et al., 2018).

Chlamydomonas *ape1* mutants show an increased light sensitivity compared to WT, due to the ROS formation as indicated by increase in GPX5 (Figure 7, (Fischer et al., 2009; Roach et al., 2017)). The increase in ROS, more specifically ^1^O_2_, results from destabilized PSII/impaired acclimation but this phenotype is alleviated under high CO_2_ conditions (Figure 1). It appears as a common trait that when a regulatory pathway is missing, affecting either light-harvesting, electron transport or CO_2_ capture, that growth is retarded under restrictive conditions (rather high light >300 μmol_photons_.m^-2^.s^-1^, limiting CO_2_) while more permissive conditions (either moderate light <100 μmol_photons_.m^-2^.s^-1^ or high CO_2_) restore normal growth. Other defects in regulation of photosynthesis have similar consequences on phototrophic growth: *pgr5* in *C. reinhardtii* and *Arabidopsis* (Johnson et al., 2014; Munekage et al., 2008), *pgrl1* (Dang et al., 2014), *cas* (Wang et al., 2016), *psbS* (Correa-Galvis et al., 2016b), *npq4* (Chaux et al., 2017), as well as 1 in 5 mutants annotated as *acetate requiring* in (Dent et al., 2005). Under limiting CO_2_, linear electron flow is limited at the PSI acceptor-side and in green algae O_2_ is also used as an alternative electron acceptor to relieve acute reduction of electron carriers through the Flv Pathway (Chaux et al., 2015) or through the shuttle of reduced metabolites to the mitochondrion for oxidative phosphorylation (Dang et al., 2014; Larosa et al., 2018). Whereas O_2_ photoreduction mostly damages PSI as observed in mutants impaired for cyclic electron flow (Johnson et al., 2014), an absence of APE1 results in PSII photodamage (Figure 1, 7 and Supplemental table 1).

The biophysical assay of antenna size, α- and β-centers and connectivity (Figure 8) confirmed that despite changes to supramolecular membrane organization that affect effective antenna size, the functional interaction between PSII units (the connectivity experiment to measure excitonic coupling) were not affected between *ape1* and the wild type. This suggests again for APE1 a regulatory role rather than at the level of the PSII function. Similarly, APE1 is not required for PSII biogenesis (Figure 3). Neither did we find it to be directly involved in PSII repair or *de novo* synthesis (Figure 7). Instead, an accelerated degradation of D1 protein as well as an increased accumulation of GPX5 were observed in *ape1* (Figure 7) and both these effects are linked to singlet oxygen production at the level of PSII. PSII photoinhibition increases linearly with light intensity (Tyystjarvi and Aro, 1996) and ROS production in isolated thylakoids in vitro is dependent on membrane stacking (Khatoon et al., 2009). Thus, the light sensitivity and PSII photoinhibition observed in *ape1* is correlated to the retention of PSII SC and the greater number of appressed membranes (Figure 7C and 10). The *ape1* mutant has a reduction in PSII core monomers and RC47 when grown photrophically in low light (Figure 4) more likely due to the remodeling of PSII cores into SC and not simply due to a deficient PSII repair cycle. However, as PSII heterogeneity is important for repair, it is clear that there is some overlap between these two processes. The lack of coordination between PSII core and antenna under non-stressful conditions, that results in suboptimal accumulation of PSII core proteins, D1 and D2 (−20%) and an increase in total LHCII antenna (+20%) (Figure 3) suggests that APE1 has a constitutive function in maintenance of PSII, in line with a regulatory role in light acclimation.

When PSII is photodamaged, D1 requires repair and the first step is the disassembly of PSII SC to core monomers followed by unfolding of thylakoid membranes (Lu, 2016; Theis and Schroda, 2016). Psb29 / Thylakoid Formation 1 (THF1) is involved in this disassembly process via interaction with the FtsH protease and LHCII antenna (Bec Kova et al., 2017; Huang et al., 2013). In our model, APE1 responds to an increase in light by releasing the oligomeric structure of PSII SC, prior to the disassembly of PSII megacomplexes by THF1 and FtsH protease for PSII repair illustrated by the model (Figure 11). APE1 binds to PSII core and not LHCII, perhaps preferentially accumulated in stromal lamellae and margins of stacked regions as measured in (Tomizioli et al., 2014). When in PSII SC, it releases core complexes as dimers and monomers. As a consequence, PSII heterogeneity influences the stacking of Chlamydomonas thylakoids, promoting longer and less stacked membranes with an increased volume of end membranes.

**Figure 11.**
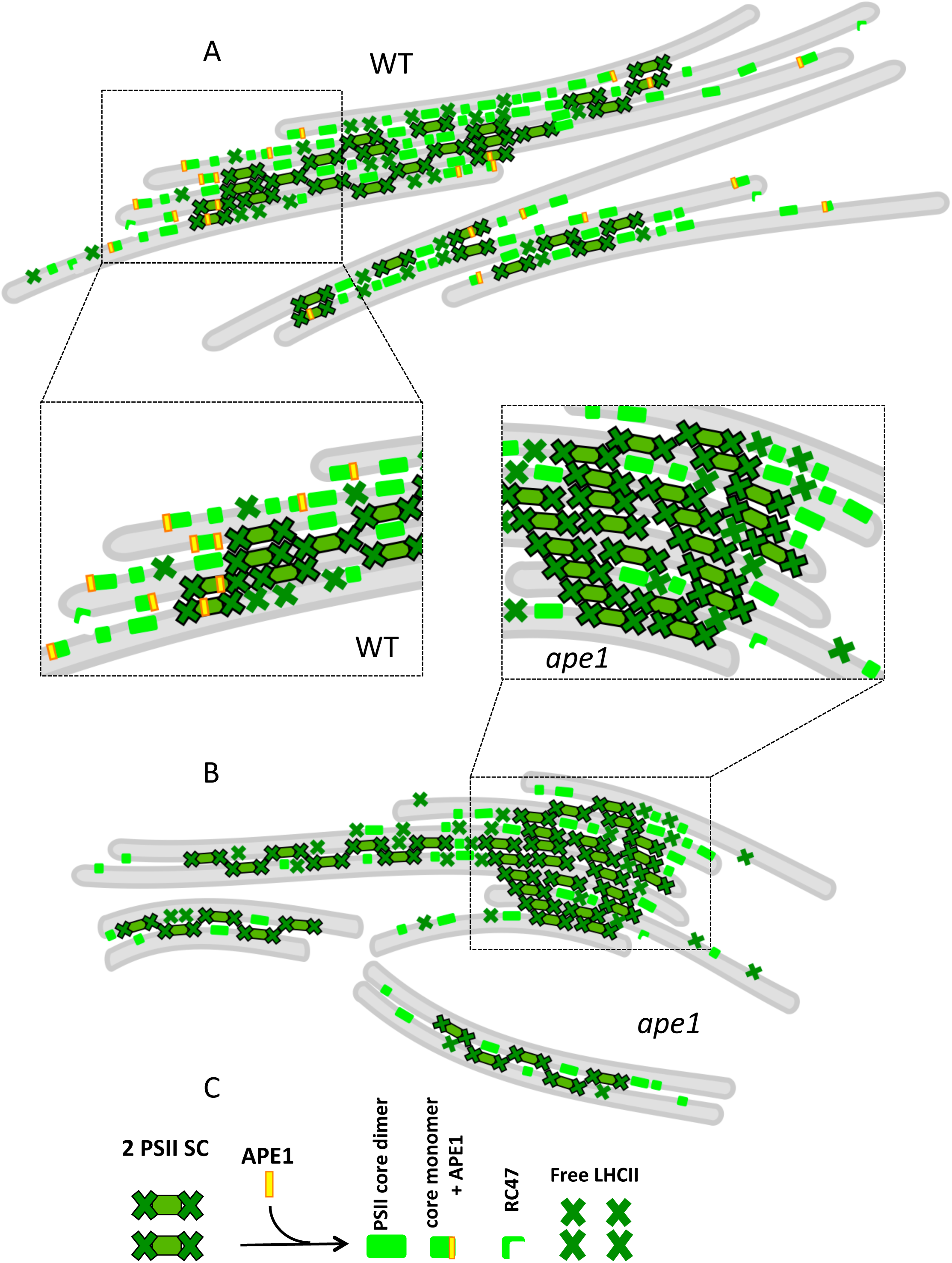
**Model: APE1 releases PSII supercomplexes in *Chlamydomonas reinhardtii*.** Thylakoid membranes (grey) and lumen (pale grey) harbor PSII of different oligomeric states. (To focus on PSII, other photosynthetic complexes are omitted in this illustration.) **A.** APE1 is bound to PSII core and not LHCII, when in PSII supercomplexes it releases core complexes. PSII complexes are heterogeneous in their oligomeric composition in cells grown phototrophically under ambient CO_2_. As a consequence, PSII plasticity influences the stacking of Chlamydomonas thylakoids. **B.** In the absence of APE1, PSII supercomplexes are more stable and interactions between the LHCII influence greater stacking of membranes that lead to a reduction in end-membranes and non-apressed membranes. It also affects the number of PSII cores that are free and not bound to LHCII and the total accumulation of PSII. **C.** An example equation for APE1 function: two PSII complexes plus interaction with APE1 results in PSII core dimer, PSII core monomer with APE1, RC47 + free LHCII

Worth commenting on is the effect on PSI complexes observed in *ape1-2* (Figures 4,7 and 9). After ruling out an interaction with PSI (Figure 5) we propose that the more compact and more stacked membranes in *ape1-2* compared to the wild type likely result in greater stability and fractionation of the higher molecular mass PSI-LHCA complexes. This was similarly observed in the *PsbN* mutant, a *bona fide* PSII repair protein, with as a consequence disrupted thylakoids, that accumulates higher order PSI complexes (Torabi et al., 2014).

During high light acclimation in plants, it takes up to 2 days for the antenna proteins to be degraded. Yang and co-workers (Yang et al., 1998) found that none of the known chloroplast proteases were involved in this process. This suggests the existence of a new unknown process in the degradation of antennae that could be autophagy. In a study of ATl1t (Autophagy-related 8 interacting protein 1) using split ubiquitin and BiFC assays (Michaeli et al., 2014), ATL1 was shown to interact with 13 chloroplast proteins including APE1 and PsbS. The exact interpretation of this result is unclear, but it could suggest that photoprotection proteins, which are necessary under stress conditions, are resistant to proteases and are only targeted for degradation under conditions where cell death is impending. It could also point to a more complex role for APE1 and PsbS as markers for selective high light acclimation as vesicle cargo for plastid remodeling (Baena-Gonzalez and Sheen, 2008; Khan et al., 2013).

Very few specific regulators have been directly attributed to the light acclimation process since it was first described in Björkman’s pioneering work (Björkman and Ludlow, 1972). Photosystem and antenna remodeling in response to light intensity characterized there and in later studies (Anderson et al., 1988; Melis, 1991) is coherent with what is witnessed in the *ape1* mutant phenotype of both Chlamydomonas and Arabidopsis (Walters et al., 2003). We envisage two possibilities for a mode of action: (1) Via an interaction with the PSII core, APE1 may destabilize the density packing of thylakoids by impairing hydrophobic interactions between transmembrane helices. (2) APE1 contains a soluble domain, with highly conserved charged residues (Glu, Asp, Arg, His) (shown in Supplementary Figure 2) which may bind, release or sense a substrate that changes electrostatic or Van der Waals attraction between polypeptides across the stromal gap. We predict that APE1 may only be involved in a very initial stage of the light acclimation process whereby it responds to light by enabling PSII SC to disassemble promoting PSII core complex heterogeneity and thylakoid destacking. Only under high light conditions, in Chlamydomonas, does this process become critical for survival.

## Materials and Methods

### Strains

The wild type strain (Jex4 *mt^-^*) used in this study is a progeny of 137c backcrosses. *ΔrbcL 2A mt^-^* strain used as the recipient strain for generation of the mutant library comes from a backcross of *ΔrbcL* strain (Johnson et al., 2010) and described as the “control” strain in figure 1. *ΔrbcL 2A mt-* strain was transformed with the *aphVIII* cassette obtained by digestion with SacI and KpnI of the pBC1 plasmid derived from pSI103 (Sizova et al., 2001). Cells were plated on TAP containing 15 μg.mL^-1^ Paromomycin. *ape1-1* strain is the progeny of *ΔrbcL ape1* obtained in the generated mutant library, outcrossed to Jex4 *mt+* twice*. ape1-2* strain is the progeny of two outcrosses of strain LMJ.RY0402.039504 from CliP (Li et al., 2016) with Jex4 *mt^+^* and *mt^-^.* Complemented strains were generated as described in “Transformation” using the vector obtained as described in “vector construction for nuclear complementation” below. *psbA* and *psaB* are the Fud7 and C3 mutants, a gift from F.A. Wollman.

### Cell culture

Cell cultures were grown in Tris-Acetate-Phosphate medium at 25°C under ambient air at 10 μmol_photons_.m^-2^ s^-1^ in incubation shakers, this is referred to as heterotrophic conditions in the text, and when stated, shifted to phototrophic conditions by centrifugation and resuspension in minimal media under ambient air. Cells were always kept at exponential phase by daily dilution to a density of around 1-2.10^6^ cells.mL^-1^. Experiments performed using batch culture were done in 250 or 500 mL Erlenmeyer flasks. Standard recipes were used as in (Harris, 1989). Maintained cultures were always cultivated at < 10 μmol_photons_.m^-2^ s^-1^ in solid TAP media.

### Turbidostats

Photobioreactors were run as turbidostats (OD_880_ maintained at 0.4 by addition of fresh minimal medium). Set-up was the same as in (Chaux et al., 2017), except that the CO_2_ input was maintained constant at ambient level (correction to avoid natural daily CO_2_ variation). Light conditions used were 40 and 320 μmol_photons_.m^-2^.s^-1^. Samples were taken by pressurizing the culture tank, for microscopy analysis, BN-PAGE, pigment content, and photosynthetic parameters. Wild type and *ape1-2* cells were grown in triplicates in photobioreactors in low light until their growth in these conditions stabilized, we called this an “acclimated state”. We took samples of these cultures for detailed biochemical analysis. Cells that had reached an acclimated state at low light were then subjected to high light, saturating for photosynthesis. Again, we allowed the growth rates of the cells to stabilize (about 5 days) before sampling the cultures.

### DNA extraction

DNA extraction was performed either according to a phenol-chloroform method or according to a rapid total DNA extraction with Chelex 100 (Sigma).

### RESDA-PCR adapted from (Gonzalez-Ballester et al., 2005)

First amplification was done in 20 μL with DegPstI (degenerated primers) and Rb1 (on the *aphVIII* cassette) primers with Taq polymerase (NEB) using 58°C annealing temperature for 5 cycles, *25°C and then 55°C for 1 cycle, 58°C for 2 cycles, 40°C for 1 cycle*, repeated 20 times between *. The PCR mix was then diluted 1000 times and 1 μL used as matrix for the second amplification. The latter was done using KOD Xtreme polymerase (Novagen) with Q0 (on an adaptator on DegPstI primer) and Rb4 (on the cassette), using 60°C annealing temperature for 32 cycles. Fragments were loaded on 1% agarose gels, the band of interest excised from the gel and cleaned using Macherey-Nagel™ NucleoSpin™ Gel and PCR Clean-up Kit and cloned into pGEM-T (Promega) for amplification before sequencing.

### Genome mapping

*ΔrbcL ape1* full insertion site in *APE1* was amplified from phenol-chloroform extracted DNA using primer A (5’-CATAACCCAGACCCCCAGAG-3’) and B (5’-CAGACTCATCCGGACCCCAA-3’) on genomic DNA, from either side of the insertion, using LA-Taq (Thermo Scientific, 60°C, 30 cycles). LMJ.RY0402.039504 was subcloned, and DNA extracted using the Chelex 100 method. The 3’ side of the insertion was amplified using primer D (5’-GACGTTACAGCACACCCTTG-3’) on the cassette and C (5’-ATCTCTTTCGGGTCCATCCT-3’) primer 600 bp upstream on genomic DNA (LA-Taq, 60°C, 30 cycles).

### RT-PCR

Cells were grown in TAP low light conditions. RNA was isolated using the PureLink RNA reagent from Invitrogen according to the manufacturer instructions, treated by TURBO DNAse enzyme (Applied Biosystems) and purified with NucleoSpin RNA Clean-up kit (Macherey-Nagel). RNA was quantified by Nanodrop (Thermoscientific). cDNA was synthesised using the standard SuperScript III protocol (Invitrogen) with oligo-dT primers. cDNA was amplified by PCR using the housekeeping gene, *RAK* as positive control and using primers E (5’-ACCTATCCTCGCTGTTCCTG-3’) and F (5’-CACGTCCTTCTTGTCCTTGC-3’) for *APE1* for 30 cycles.

### Genetic transformation

300 μL of cell culture resuspended at around 1-2.10^8^ cells.mL^-1^ in TAP 60 mM Sucrose were aliquoted in 0.4 cm gapped cuvettes. 1 μg of linearized DNA and 4 μL of salmon sperm DNA were added and cells were incubated 20 min on ice. Negative controls were done without adding DNA. Electroporation conditions were 1000 V, 25 μF. Cells were allowed to recover in low light overnight in 10 mL TAP 60 mM Sucrose, centrifuged and resuspended in 500 μL TAP. Volumes from 50 μL to 200 μL were plated on TAP containing antibiotics to obtain an optimal colony density. The *ape1-2* mutant was transformed with a plasmid containing Zeo^R^ cassette and the genomic version of the wild type *APE1* under its native promoter (1500 bp upstream of the start site of *APE1*). Transformants were selected on zeocin-containing plates and were screened using chlorophyll fluorescence imaging

### Vector construction for nuclear complementation

Transformation plasmid was constructed using restriction site cloning in pMS188 (Schroda et al., 2002) containing a bleomycin resistance cassette. *APE1* gene with an additional 1500 bp on the 5’ side and 500 bp on the 3’ side was amplified by PCR using Q5 High-Fidelity DNA Polymerase (NEB) using primers 5’-<u>TTATAA</u>TACCCACCCGTCAAAGCTGTG-3’ and 5’-<u>TTATAA</u>CAGACTCATCCGGACCCCAA-3’ containing an additional <u>PsiI restriction site</u> (70°C, 30 cycles). pMS188 was digested using PsiI and dephosphorylated. Ligation was done in 20 μL overnight at 4°C using 50 ng of pMS188 and 140 ng of the purified PCR product. DH5-α bacterial strains were transformed and plated on plates containing Kanamycin. Presence and orientation of *APE1* gene in pMS188 was then checked by digestion using XbaI, yielding 3 fragments of 1.6, 2.9 and 4.5 kb on 1% agarose gel. The construct was then verified by sequencing using 5’-TACCCACCCGTCAAAGCTGTG-3’, 5’-GGCGTCTTCCACACTCACTG-3’, 5’-CACACCACTCCCGTAGCTGA-3’, 5’-GAGGTGATCGCTTCGGTAGG-3’ and 5’-TGGACGCAAATGGAAACAAG-3’ primers. The transformation plasmid was then linearized with ScaI-HF, transformation was done as stated above, and cells were plated on TAP containing Zeocin at 45 μg.mL^-1^.

### Recombinant APE1 protein production

*APE1* cDNA without the sequence coding for the transit peptide (APE1 total protein, 223 amino acids) or without the sequence coding for the transmembrane domain (APE1 soluble region, 156 amino acids) was introduced in pLIC03 vector (LIC : ligation-independent cloning; (Aslanidis and de Jong, 1990) using GoldenGate cloning technique. The pLIC03 vector is the pET-28a+ vector (Novagen) modified to add downstream of the start codon a 6×His tag and a TEV protease-cleavage site. These are followed by the suicide gene *sacB* flanked by *BsaI* restriction sites, replaced by *APE1* cDNA flanked by complementary *BsaI* restriction sites sequences. 50 ng of vector was used with 1:3 cDNA for six successive rounds of digestion (*BsaI*, 37°C, 5 min) and ligation (T4 ligase, 16°C, 10 min) in the same mix. Successful ligation of *APE1* cDNA removes the *BsaI* recognition sequence. The cycles were ended by a final ligation step (30 min at *E. coli* cells were then transformed with the ligation products for vector amplification, screening and sequencing. 65 ng of vectors were then used to transform Rosetta *E. coli* cells cultured in TB medium at 37 °C up to OD 1. IPTG was added to induce APE1 recombinant protein expression, temperature was decreased to 17 °C and the cells were grown for an additional 18 h. Cells were harvested by centrifugation and the pellet was resuspended in lysis buffer during 30 min at 4°C. Lysis buffer contained 300 mM NaCl, 50 mM Tris pH 8.0, 10 mM imidazole, 5% (w/v) glycerol, 0.25 mg.mL^-1^ lysozyme, 0.1% Triton, 1mM EDTA and protease inhibitors. Cells were lysed by sonication, and incubated at 25°C with 20 mM MgSO_4_ and 10 μg.mL^-1^ DNase. The lysate was then centrifuged; the supernatant was collected and incubated for 5 min on 1 mL Ni Sepharose^®^ 6Fast Flow (GE Healthcare) equilibrated with 10 mL binding buffer. The resin was washed (5 x 1mL) and eluted (5 x 1mL). Buffers contained 300 mM NaCl, 50 mM Tris pH 8.0, 5% (w/v) glycerol; with 10 mM imidazole for binding, 50 mM for washing and 250 mM for elution.

### Antibody generation

APE1 recombinant protein containing only the soluble part was sent to ProteoGenix for antibody production from rabbit. 9 mL of filtered immune serum were purified against the soluble part of the recombinant protein (10 mg) covalently coupled to 1mL HiTrap NHS-activated HP resin (GE Healthcare) according to the manufacturer instructions. The serum recirculated on the resin for 2 hours at a flow rate of 0.5 mL/min and was eluted at pH 3. The eluted fraction was collected in neutralizing buffer. The antibody is used at 1:10000 dilution for immunoblotting.

### Spot tests

25μL of cells at 10^6^ cells.mL^-1^ were spotted on TAP or MIN plates, allowed to dry and placed at different light intensities as shown for 5-14 days.

### Pigment quantification

Chlorophyll content was measured in acetone 80% according to Porra et al. (1989) or in methanol using:

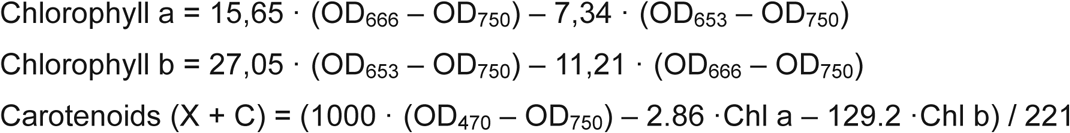

### Protein Analysis

Proteins were extracted from total cells or thylakoids with cold acetone 80% and separated under denaturing conditions on 13% SDS-PAGE gels, unless otherwise indicated. Proteins were loaded based on chlorophyll content (1 μg), and total protein amount was quantified on Coomassie Blue gels to adjust the loading. SYPRO Ruby Protein gel stain was performed according to manufacturers protocol (Molecular Probes, Invitrogen). Proteins were transferred onto nitrocellulose membranes (BioTrace NT, Pall Corporation) using liquid transfer (except for Figure 4D where 10% NativePAGE Bis-Tris MES gel (Invitrogen) and a semi-dry transfer was performed. All primary antibodies used were sourced from Agrisera. APE1 antibody was generated in a rabbit against the soluble part of the recombinant protein. Secondary antibodies used were always HRP-conjugated anti-rabbit (Invitrogen). HRP-peroxidase chemiluminescent substrate (Invitrogen) was used to reveal the antibody signal using the GBOX imaging system (Syngene).

### Thylakoid extraction and native protein analysis

Thylakoid extraction was done according to the standard method (Chua and Bennoun, 1975). Thylakoids were resuspended in Hepes 5 mM, EDTA 10 mM at 1 mg/mL chlorophyll for subsequent SDS-PAGE analysis. For density gradient analysis and crosslinking, 1% digitonin (final) and then 1% n-Dodecyl- α -D-Maltoside (final) with 0.35% glutaraldhyde was loaded onto sucrose gradients and treated as in (Caffari et al.,ref). For non-denaturing conditions, thylakoids were resuspended in NativePAGE sample buffer (Life technologies) at 1 mg.mL-1 chlorophyll, thylakoids were solubilized for 5 min on ice in the same volume of 2% n-Dodecyl-β-D-Maltoside (0.5 mg.mL-1 chlorophyll and 1% n-Dodecyl-β-D-Maltoside final). Differential solubilization was achieved by using 1% digitonin (final) and then 1% n-Dodecyl-β-D-Maltoside (final) on the remaining non-solubilized material. For interaction analysis, thylakoids were solubilized with 0.5% digitonin (final) and 0.5% n-Dodecyl-α-D-Maltoside (final). Concentrations and resulting profiles were similar to that observed in (Pagliano et al., 2012) and were confirmed in figure 6. For each analysis, 20 μL were then loaded with 2 μL of G-250 sample additive (Life technologies) on 4-16% (Figures 4A and 7) or 3-12% (Figures 4C and 6) NativePAGE gels (Life technologies). Cathode Running buffer (Life technologies) was supplemented with 0.02% G-250 for 2/3 of the migration, and with 0.002% G-250 for the remaining third. For second dimension analysis, bands were incubated 1h at room temperature in LDS, 50 mM DTT and 5 M Urea and loaded on 13% 5M urea SDS-PAGE. Gels were then either transferred on nitrocellulose or silver stained. Proteins separated by BN-PAGE were identified using immunoblots and silver staining on the second dimension, and by comparison to similar results in the literature (Drop et al., 2011; Drop et al., 2014; Muranaka et al., 2016; Rexroth et al., 2003). Spectra of gradient fractions (350 - 750 nm) were obtained using UV-Vis spectrophotometer (Varian Cary 300) at a scan rate of 240 nm/min and baseline correction.

## Chlorophyll Fluorescence Analysis

Fluorescence measurements on Petri dishes were performed using the set up for *in vivo* chlorophyll fluorescence imaging (Beal SpeedZen Camera) (Johnson et al., 2009). Pulse-Amplitude-Modulation Fluorimeter (Walz) was used for chlorophyll fluorescence kinetics and monitoring of photoinhibition. Dark-adapted cells were subjected to a saturating pulse (8 000 μmol_photons_.m^-2^.s^-1^) to measure F_v_/F_M_ or exposed to a given light intensity (red light) and Φ_PSII_ was probed every minute. Using the saturating pulse but with fast sampling kinetics, OJIP was monitored over 300 milliseconds. Heterogeneity of PSII centres and connectivity of PSII centres was performed using the Joliot-type Spectrophotometer (JTS-Biologic). Melis has shown that the complementary area over the fluorescence curve (proportional to the reduction of Q_A_) of DCMU poisoned sample shows two phases in a semi-logarithmic plot (Melis and Homann, 1976). The fast phase, or α-centers, appearing as a straight line at 0 < t < 50 ms corresponds to PSII with a large light harvesting capacity, whereas the slow phase, or β-centers, visible between 100 < t < 250 ms corresponds to PSII centers with less associated chlorophylls (core complex). The intercept of the slow component at t = 0 (dotted lines) allows for quantification of the α-centers. Connectivity was performed as in (Cuni et al., 2004) 10 μM DCMU was used to block electron transfer beyond Q_A_. Connectivity between PSII centers is illustrated by the non-linearity between the variable part of chlorophyll fluorescence yield (probability to reemit a photon) against the relative concentration of reduced Q_A_ and data were fitted to *F*_V_ = [Q_A_^-^] / (1 + J - J [Q_A_^-^]) yielding the connectivity parameter J.

### Photoinhibition experiments

Cells were grown in TAP media at 10 μmol_photons_.m^- 2^.s^-1^, diluted to a similar chlorophyll amount (3 μg.mL^-1^), allowed to stabilise for 1 hour at 40 μmol_photons_.m^-2^.s^-1^, and then transferred to 1800 μmol_photons_.m^-2^.s^-1^ light for 1 hour with or without 0.1 mg/mL chloramphenicol and 0.5 mg/mL lincomycin. Recovery was done at 40 μmol_photons_.m^-2^.s^-1^. *F*_V_/*F*_M_ was used as a non-invasive measure of the maximum PSII quantum efficiency to monitor the degree of photoinhibition caused by the light treatment. Cells were placed in the dark for 3 minutes with agitation before the measurement by PAM.

### Electronic Transmission Microscopy

This experiment was performed twice on two different sets of cultures: cells were grown phototrophically and samples were taken either from turbidostats grown at 40 μmol_photons_.m^-2^.s^-1^ or from batch culture flasks grown at 80 μmol_photons_.m^-2^.s^-1^ (statistics are a mix of both samples). Cells were collected by centrifugation and fixed with 2.5% glutaraldehyde in 0.1 M sodium cacodylate buffer, pH 7.4, at 4 °C for two days. They were then washed three times using the same buffer. Samples were post-osmicated with 1% osmium tetroxyde in cacodylate buffer for 1 h, dehydrated through a graded ethanol series, and finally embedded in monomeric resin Epon 812. All chemicals used for histological preparation were purchased from Electron Microscopy Sciences (Hatfield, USA). 90 nm ultrathin sections for transmission electron microscope (TEM) were obtained by an ultramicrotome UCT (Leica Microsystems GmbH, Wetzlar, Germany) and mounted on copper grids. They were examined in a Tecnai G^2^ Biotwin Electron Microscope (ThermoFisher Scientific FEI, Eindhoven, the Netherlands) using an accelerating voltage of 100 kV and equipped with a CCD camera Megaview III (Olympus Soft imaging Solutions GmbH, Münster, Germany). For each replicate, several photographs of entire cells and at least 20 micrographs of local detailed structures were taken, analyzed and compared.

## Accession numbers

Sequence data for *C. reinhardtii APE1* from this article can be found in GenBank under the accession number NW_001843882.1.

## Supplementary dMaterial files

**Fig S1.** Mapping of the insertion in the *APE1* gene of Chlamydomonas in two alleles.

**Fig S2.** Summary scheme of highly conserved domains in APE1: Alignments of predicted protein sequences for *APE1* gene products.

**Fig S3.** Silver stain of 2^nd^ dimension BN-PAGE of thylakoid proteins of *ape1-2*, *psbA* and *psaB* cells treated by high light in batch cultures.

**Fig S4.** Silver stain of 2^nd^ dimension BN-PAGE thylakoid proteins of WT and *ape1-2* acclimated to high light in photobioreactors run as turbidostats.

**Fig S5.** TEM images at two different magnifications of thylakoid membranes from wild type and *ape1-2*.

**Table S1.** Photosynthetic parameters in different conditions

**Table S2.** Dimensions and membrane surfaces of the grana (stacked regions) in WT and *ape1-2* cells.

**Table S3.** List of cyanobacterial proteins conserved across all oxygenic phototrophs having the same genomic co-occurrence as APE1

## Acknowledgements

We would like to thank Jerome Lavergne for invigorating and informative discussions and Corinne Cassier-Chauvat for help on cyanobacterial genome analysis. We acknowledge and are thankful to Stéphanie Blangy, Audrey Beyly-Adriano, Pascaline Auroy and Véronique Cardettini for technical assistance. This work was supported by a grant from the Agence National pour la Recherche (ChloroPaths : ANR-14-CE05-0041-01). M.C. was supported by a scholarship from the Ministery of Science and Education.

## Author Contributions

MC, JA, JD, MF, GP, BG, XJ designed the research; MC, JA, SC, PB, SC, JD, MF, XJ performed research; MC, JA, SC, PB, SC, JD, MF, GP, BG, XJ analyzed data; and MC, JA, SC, XJ wrote the paper.

